# Assessing anaerobic gut fungal (Neocalliamstigomycota) diversity using PacBio D1/D2 LSU rRNA amplicon sequencing and multi-year isolation

**DOI:** 10.1101/2020.03.24.005967

**Authors:** Radwa A. Hanafy, Britny Johnson, Noha H. Youssef, Mostafa S. Elshahed

## Abstract

The anaerobic gut fungi (AGF, Neocallimastigomycota) reside in the alimentary tracts of herbivores where they play a central role in the breakdown of ingested plant material. Accurate assessment of AGF diversity has been hampered by inherent deficiencies of the internal transcribed spacer1 (ITS1) region as a phylogenetic marker. Here, we report on the development and implementation of the D1/D2 region of the large ribosomal subunit (D1/D2 LSU) as a novel marker for assessing AGF diversity in culture-independent surveys. Sequencing a 1.4-1.5 Kbp amplicon encompassing the ITS1-5.8S rRNA-ITS2-D1/D2 LSU region in the ribosomal RNA locus from fungal strains and environmental samples generated a reference D1/D2 LSU database for all cultured AGF genera, as well as the majority of candidate genera encountered in prior ITS1-based diversity surveys. Subsequently, a D1/D2 LSU-based diversity survey using long read PacBio SMRT sequencing technology was conducted on fecal samples from 21 wild and domesticated herbivores. Twenty-eight genera and candidate genera were identified in the 17.7 K sequences obtained, including multiple novel lineages that were predominantly, but not exclusively, identified in wild herbivores. Association between certain AGF genera and animal lifestyles, or animal host family was observed. Finally, to address the current paucity of AGF isolates, concurrent isolation efforts utilizing multiple approaches to maximize recovery yielded 216 isolates belonging to twelve different genera, several of which have no prior cultured-representatives. Our results establish the utility of D1/D2 LSU and PacBio sequencing for AGF diversity surveys, and the culturability of a wide range of AGF taxa, and demonstrate that wild herbivores represent a yet-untapped reservoir of AGF diversity.

## Introduction

Members of the anaerobic gut fungi (AGF) are strict anaerobes that inhabit the rumen and alimentary tract of a wide range of foregut and hindgut herbivores. The AGF play an important role in the breakdown of ingested plant biomass via enzymatic and physical disruption in the herbivorous gut^1^. AGF represent a distinct basal fungal phylum (Neocallimastigomycota) that evolved 66 (±10) million years ago coinciding, and possibly enabling, mammalian transition from insectivory to herbivory^2^.

Culture independent amplicon-based diversity surveys have been widely utilized to gauge anaerobic fungal diversity and community structure in herbivores^3, 4, 5, 6^. The internal transcribed spacer1 (ITS1) region within the ribosomal operon has been almost exclusively utilized as the phylogenetic marker of choice in culture-independent sequence-based phylogenetic assessments of AGF diversity^7^ Such choice is a reflection of its wider popularity as a marker within the kingdom Mycota^8, 9^, the high sequence similarity and limited discriminatory power of the 18S rRNA gene between various AGF taxa^10^, and its relatively shorter length, allowing high throughput pyrosequencing- and Illumina-based diversity assessments^3, 6^. However, concerns for the use of ITS1 in diversity assessment for the Mycota^11^, basal fungi^12^, and the Neocallimastigomycota^7^ have been voiced. The ITS1 region is polymorphic, exhibiting considerable secondary structure (number and organization of helices^13^), and length^14^ variability. Such polymorphism renders automated alignments, reproducible sequence divergence estimates, and classification of sequence data unreliable and highly dependent on alignment strategies and parameters specified. In addition, significant sequence divergence between copies of the ITS1 region within a single strain have been reported (up to 12.9% in^15^), values that exceed cutoffs utilized for species (even genus in some instances) level delineation from sequence data^3, 16, 17, 18^. Such limitations often necessitate laborious subjective manual curation and secondary structure incorporation into alignment strategies^13^, although it is well recognized that these efforts only partially alleviate, rather than completely address, such fundamental limitations.

The 28S large ribosomal subunit (LSU) is one of the original genes proposed for fungal barcoding^12^. Hypervariable domains 1 and 2^19^ within the LSU molecule (D1/D2 LSU) have previously been utilized for differentiating strains of AGF via molecular typing^20, 21, 22^, or sequencing^23, 24^ Compared to ITS1, D1/D2 LSU region exhibits much lower levels of length heterogeneity and intra-strain sequence divergence in fungi^25^, including the AGF^20^. Identification and taxonomic assignment of AGF strains based on D1/D2 LSU have gathered momentum; and D1/D2 LSU-based phylogenetic analysis has been reported in all manuscripts describing novel taxa since 2015^15, 26, 27, 28, 29, 30, 50^. The potential use of D1/D2 LSU as a marker in culture-independent AGF diversity surveys has been proposed as a logical alternative for ITS1^7, 14^ The lack of specific AGF primers and the relatively large size of the region (approximately 750 bp) has been viewed as a barrier to the wide utilization of short read, high-throughput, Illumina-based amplicon sequencing in such surveys. However, the recent development of AGF LSU-specific primers^24, 31^, as well as the standardization and adoption of PacBio long-read sequencing for amplicon-based diversity surveys^32, 33^ could enable this process.

Theoretically, a comprehensive assessment of diversity and community structure of a host-associated lineage necessitates sampling all (or the majority) of hosts reported to harbor such lineage. However, to date, the majority of AGF diversity surveys conducted have targeted a few domesticated herbivores, e.g. cows, sheep, and goats^4, 5, 34^. “Exotic” animals have been sampled from zoo settings only sporadically, and on an opportunistic basis^3, 35^.

Isolation of AGF taxa enables taxonomic, metabolic, physiological, and ultrastructural characterization of individual taxa. As well, cultures availability enables subsequent –omics, synthetic and system-biology, and biogeography-based investigations^36, 37, 38, 39, 40, 41, 42^, as well as evaluation of evolutionary processes underpinning speciation in the AGF^2, 43^. However, efforts to isolate and maintain AGF strains have lagged behind their aerobic counterparts mainly due to their strict anaerobic nature and the lack of reliable long-term storage procedures. Due to these difficulties, many historic isolates are no longer available, and most culture-based studies report on the isolation of a single or few strains using a single substrate/enrichment condition from one or few hosts^29, 44^. Indeed, a gap currently exists between the rate of discovery (via amplicon-based diversity surveys) and the rate of isolation of new taxa of AGF, and several yet-uncultured AGF lineages have been identified in culture-independent diversity surveys^17^. Whether yet-uncultured AGF taxa are refractory to isolation, or simply not yet cultured due to inadequate sampling and isolation efforts remains to be seen.

The current study aims to expand our understanding of the diversity of AGF while addressing all three impediments described above. First, we sought to develop D1/D2 LSU as a more robust marker for AGF diversity assessment by building a reference sequence database correlating ITS1 and D1/D2 LSU sequence data from cultured strains and environmental samples. Second, we sought to expand on AGF diversity by examining a wide range of animal hosts, including multiple previously unsampled wild herbivores. Third, we sought to demonstrate the utility of intensive sampling and utilization of various isolation strategies in recovering AGF strains and testing the hypothesis that many yet-uncultured AGF lineages are indeed amenable to cultivation. Collectively, these efforts provide an established framework for future utilization of D1/D2 LSU amplification and PacBio sequencing for AGF community assessment, highlight the value of sampling wild herbivores, and establish the culturability of a wide range of AGF taxa.

## Results

### A reference D1/D2-LSU dataset for the Neocallimastigomycota

A 1.4-1.5 Kbp amplicon product encompassing the ITS1-5.8S-ITS2-D1/D2 LSU region was amplified and sequenced from AGF pure cultures and environmental samples to correlate the D1/D2 LSU region to the corresponding ITS1 region, and to provide a reference D1/D2 database for future utilization in high-throughput diversity surveys. Using this approach, representative D1/D2 LSU of all the previously cultured AGF genera *Agriosomyces, Aklioshbomyces, Anaeromyces, Buwchfawromyces, Caecomyces, Capellomyces, Cyllamyces, Ghazallomyces, Joblinomyces, Feramyces, Khyollomyces, Liebetanzomyces, Neocallimastix, Orpinomyces, Pecoramyces, Piromyces*, and *Tahromyces* were obtained (Table 1). Representatives of the genus *Oontomyces* were not encountered in this study, but reference LSU and ITS1 sequences were obtained from prior publication^26^. In addition, representative sequences of D1/D2 LSU of candidate genera AL3, AL4, AL8, MN3, MN4, SK3, and SK4, previously identified in ITS1 culture-independent datasets were also obtained (Table 1, Datasets 1-3). Finally, representatives of six completely novel AGF candidate genera (RH1-RH6) were also identified (Table 1, Datasets 1-3) and confirmed as novel independent clades in ITS1 and D1/D2 LSU-based phylogenetic analysis. It should be noted that multiple previously reported yet-uncultured (candidate) genera have recently been successfully isolated, e.g. AL1 (*Khyollomyces*), AL5 (*Joblinomyces*), AL6 (*Feramyces*), AL7 (*Piromyces finnis*), MN1 (*Cyllamyces*), SP4 (*Liebetanzomyces*), and SK2 (*Buwchfawromyces*). In addition, some previously proposed candidate genera clustered as members of already existing genera in our analysis, e.g. SP8 with *Cyllamyces*, and SP6 with *Neocallimastix* (Table 1). As such, we estimate that only representatives of candidate genera BlackRhino, SP1, SP2^17^, and the relatively rare AL2, DA1, DT1, JH1/SP5 (ITS1 sequence representatives of these two candidate genera are 99.6% similar and so they should be considered as one candidate genus), KF1, MN2, SK1, SP3, and SP7^3, 5, 13, 17, 45, 46, 47, 48^ were not encountered in this study, and hence no reference LSU sequence data for these candidate genera are currently available (Table 1).

**Table 1.**
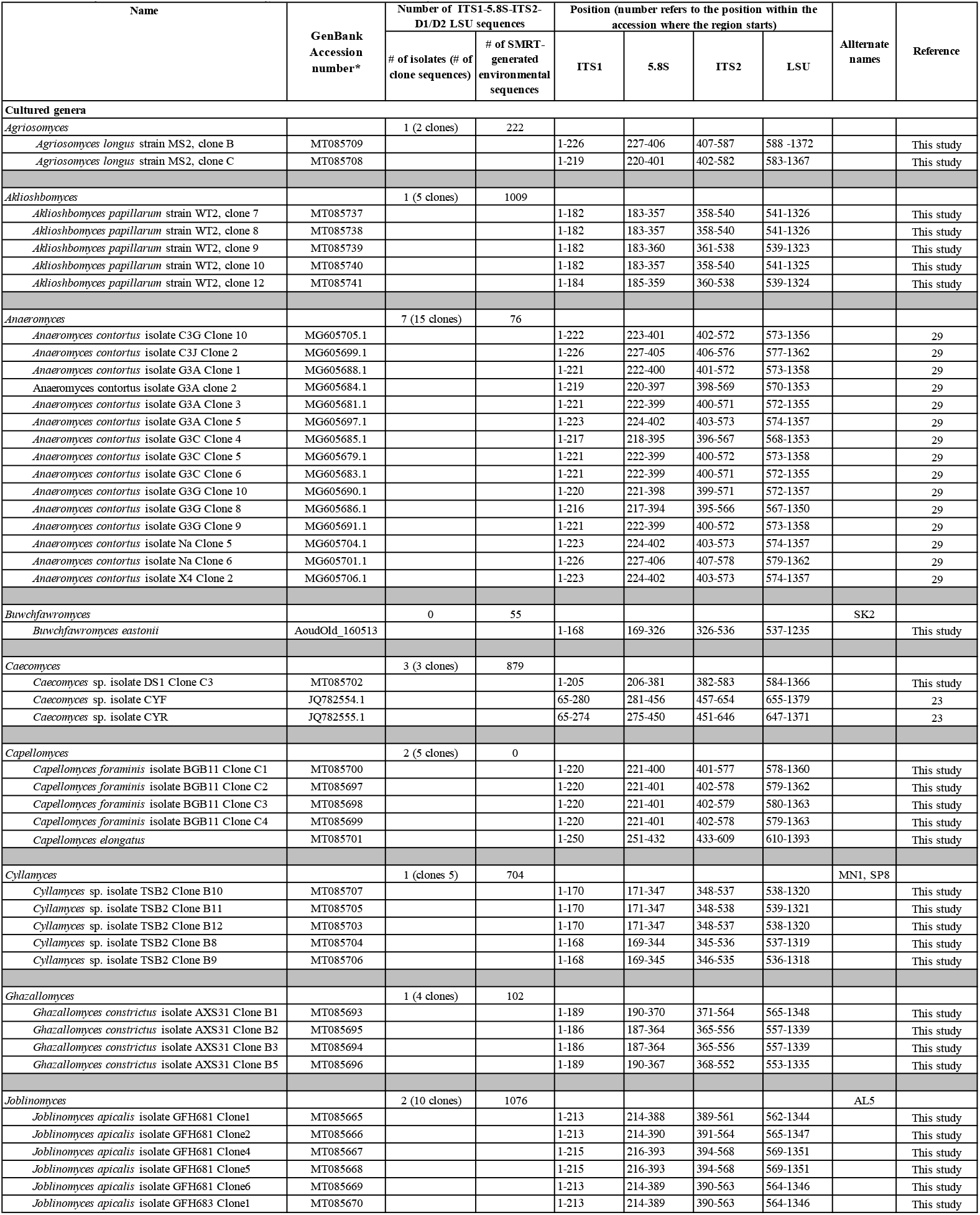
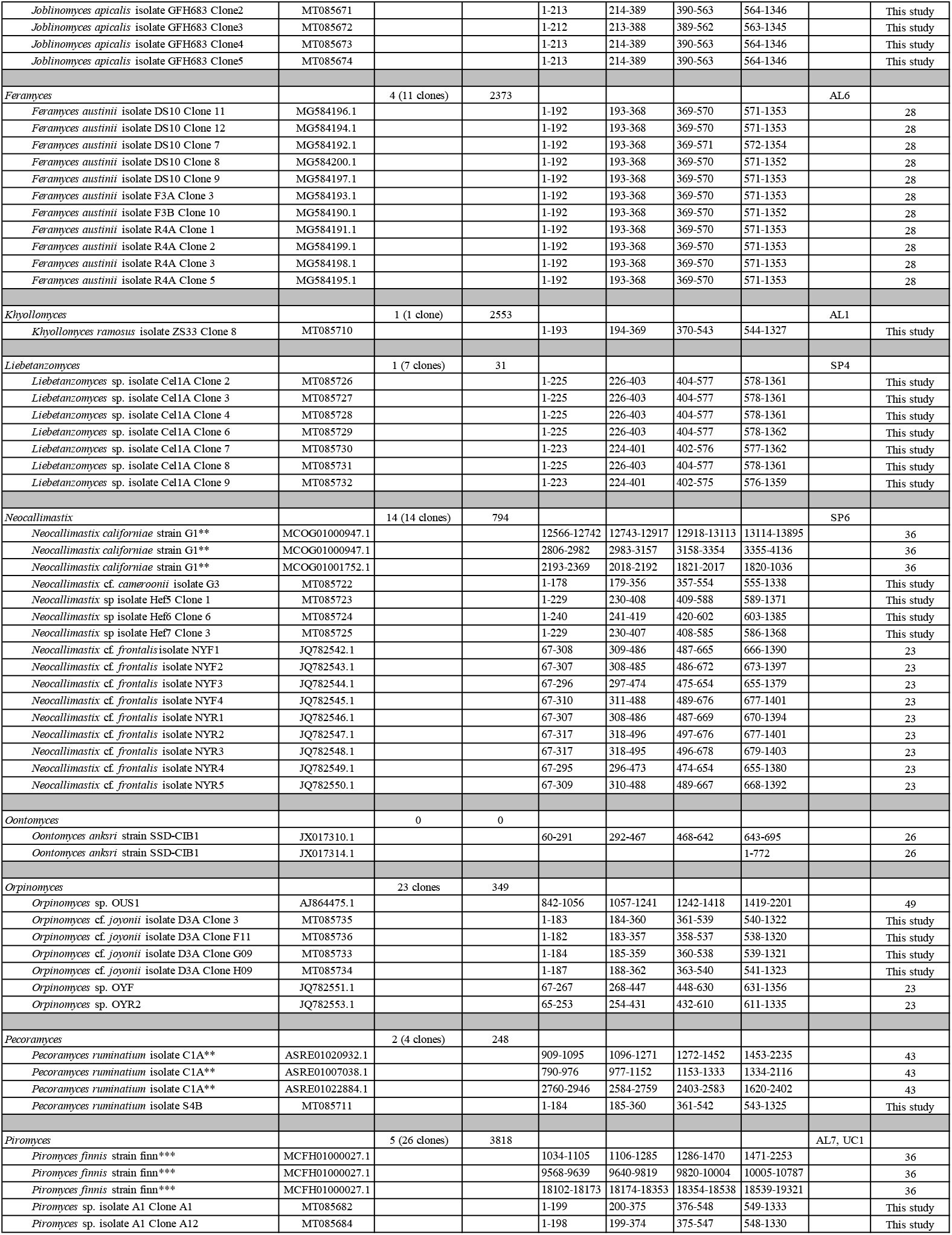
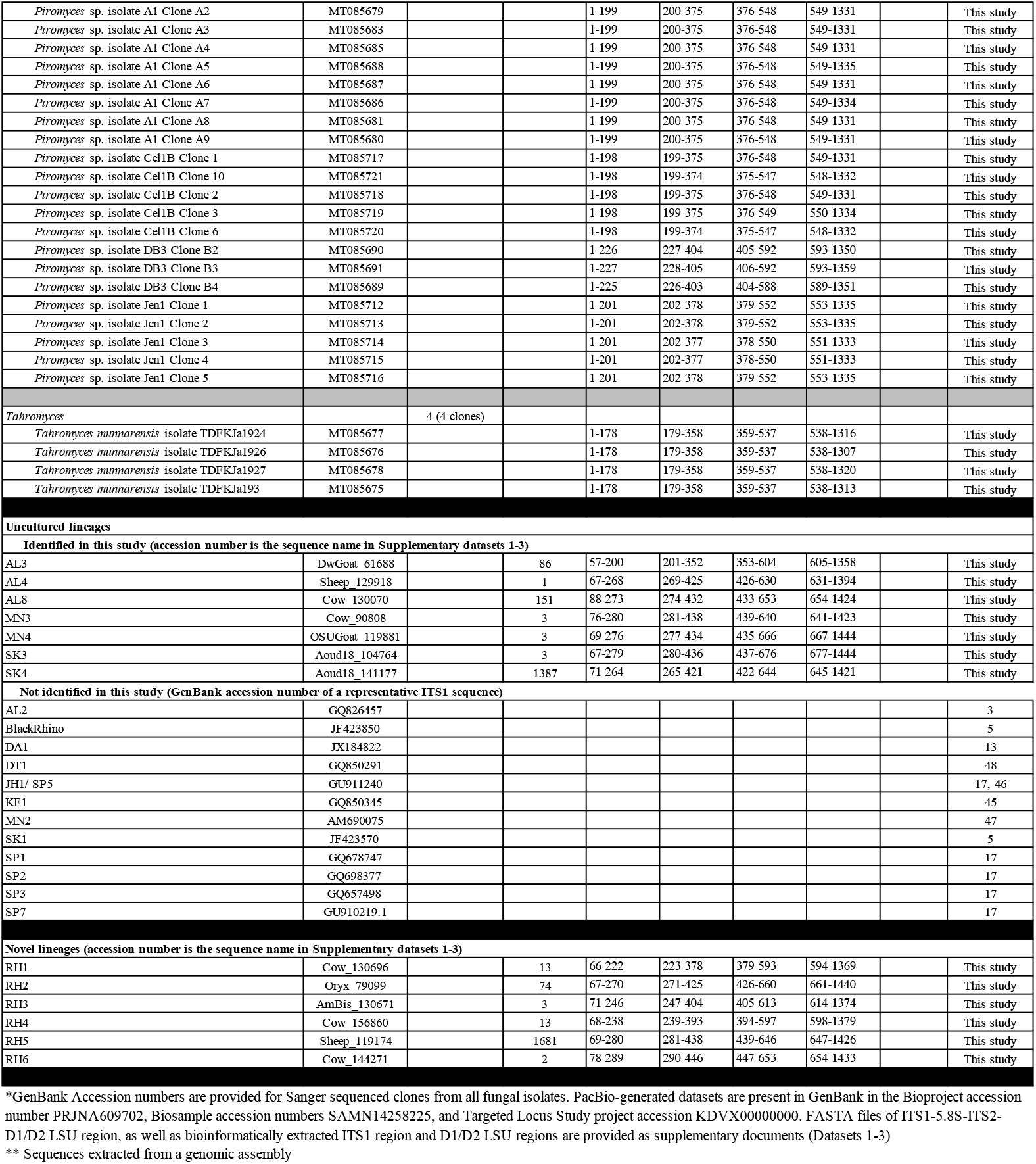
Representatives full-length sequences spanning the region “ITS1-5.8S rRNA-ITS2-D1/D2 LSU”. GenBank accession numbers are shown for all clone sequences obtained from representative AGF isolates in our culture collection. For yet-uncultured AGF taxa, accession numbers refer to the SMRT generated sequence name in Datasets 1-3. Start and end positions of ITS1, 5.8S rRNA gene, ITS2, and the D1/D2 region of the LSU are shown.

| Name | GenBank<br>Accession<br>number* | Number of ITS1-5.8S-ITS2-D1/D2 LSU sequences |  | Position (number refers to the position within the<br>accession where the region starts) |  |  |  | Allternate<br>names | Reference |
| --- | --- | --- | --- | --- | --- | --- | --- | --- | --- |
|  |  | # of isolates (# of<br>clone sequences) | # of SMRT-generated<br>environmental<br>sequences | ITS1 | 5.8S | ITS2 | LSU |  |  |
| Cultured genera |  |  |  |  |  |  |  |  |  |
| *Agriosomyces* |  | 1 (2 clones) | 222 |  |  |  |  |  |  |
| *Agriosomyces longus* strain MS2, clone B | MT085709 |  |  | 1-226 | 227-406 | 407-587 | 588-1372 |  | This study |
| *Agriosomyces longus* strain MS2, clone C | MT085708 |  |  | 1-219 | 220-401 | 402-582 | 583-1367 |  | This study |
| *Akioshbomyces* |  | 1 (5 clones) | 1009 |  |  |  |  |  |  |
| *Akioshbomyces papillarum* strain WT2, clone 7 | MT085737 |  |  | 1-182 | 183-357 | 358-540 | 541-1326 |  | This study |
| *Akioshbomyces papillarum* strain WT2, clone 8 | MT085738 |  |  | 1-182 | 183-357 | 358-540 | 541-1326 |  | This study |
| *Akioshbomyces papillarum* strain WT2, clone 9 | MT085739 |  |  | 1-182 | 183-360 | 361-538 | 539-1323 |  | This study |
| *Akioshbomyces papillarum* strain WT2, clone 10 | MT085740 |  |  | 1-182 | 183-357 | 358-540 | 541-1325 |  | This study |
| *Akioshbomyces papillarum* strain WT2, clone 12 | MT085741 |  |  | 1-184 | 185-359 | 360-538 | 539-1324 |  | This study |
| *Anaeromyces* |  | 7 (15 clones) | 76 |  |  |  |  |  |  |
| *Anaeromyces contortus* isolate C3G Clone 10 | MG605705.1 |  |  | 1-222 | 223-401 | 402-572 | 573-1356 |  | 29 |
| *Anaeromyces contortus* isolate C3J Clone 2 | MG605699.1 |  |  | 1-226 | 227-405 | 406-576 | 577-1362 |  | 29 |
| *Anaeromyces contortus* isolate G3A Clone 1 | MG605688.1 |  |  | 1-221 | 222-400 | 401-572 | 573-1358 |  | 29 |
| *Anaeromyces contortus* isolate G3A clone 2 | MG605684.1 |  |  | 1-219 | 220-397 | 398-569 | 570-1353 |  | 29 |
| *Anaeromyces contortus* isolate G3A Clone 3 | MG605681.1 |  |  | 1-221 | 222-399 | 400-571 | 572-1355 |  | 29 |
| *Anaeromyces contortus* isolate G3A Clone 5 | MG605697.1 |  |  | 1-223 | 224-402 | 403-573 | 574-1357 |  | 29 |
| *Anaeromyces contortus* isolate G3C Clone 4 | MG605685.1 |  |  | 1-217 | 218-395 | 396-567 | 568-1353 |  | 29 |
| *Anaeromyces contortus* isolate G3C Clone 5 | MG605679.1 |  |  | 1-221 | 222-399 | 400-572 | 573-1358 |  | 29 |
| *Anaeromyces contortus* isolate G3C Clone 6 | MG605683.1 |  |  | 1-221 | 222-399 | 400-571 | 572-1355 |  | 29 |
| *Anaeromyces contortus* isolate G3G Clone 10 | MG605690.1 |  |  | 1-220 | 221-398 | 399-571 | 572-1357 |  | 29 |
| *Anaeromyces contortus* isolate G3G Clone 8 | MG605686.1 |  |  | 1-216 | 217-394 | 395-566 | 567-1350 |  | 29 |
| *Anaeromyces contortus* isolate G3G Clone 9 | MG605691.1 |  |  | 1-221 | 222-399 | 400-572 | 573-1358 |  | 29 |
| *Anaeromyces contortus* isolate Na Clone 5 | MG605704.1 |  |  | 1-223 | 224-402 | 403-573 | 574-1357 |  | 29 |
| *Anaeromyces contortus* isolate Na Clone 6 | MG605701.1 |  |  | 1-226 | 227-406 | 407-578 | 579-1362 |  | 29 |
| *Anaeromyces contortus* isolate X4 Clone 2 | MG605706.1 |  |  | 1-223 | 224-402 | 403-573 | 574-1357 |  | 29 |
| *Buwchfawromyces* |  | 0 | 55 |  |  |  |  | SK2 |  |
| *Buwchfawromyces eastonii* | AoudOld_160513 |  |  | 1-168 | 169-326 | 326-536 | 537-1235 |  | This study |
| *Caecomyces* |  | 3 (3 clones) | 879 |  |  |  |  |  |  |
| *Caecomyces* sp. isolate DS1 Clone C3 | MT085702 |  |  | 1-205 | 206-381 | 382-583 | 584-1366 |  | This study |
| *Caecomyces* sp. isolate CYF | JQ782554.1 |  |  | 65-280 | 281-456 | 457-654 | 655-1379 |  | 23 |
| *Caecomyces* sp. isolate CYR | JQ782555.1 |  |  | 65-274 | 275-450 | 451-646 | 647-1371 |  | 23 |
| *Capellomyces* |  | 2 (5 clones) | 0 |  |  |  |  |  |  |
| *Capellomyces foraminis* isolate BGB11 Clone C1 | MT085700 |  |  | 1-220 | 221-400 | 401-577 | 578-1360 |  | This study |
| *Capellomyces foraminis* isolate BGB11 Clone C2 | MT085697 |  |  | 1-220 | 221-401 | 402-578 | 579-1362 |  | This study |
| *Capellomyces foraminis* isolate BGB11 Clone C3 | MT085698 |  |  | 1-220 | 221-401 | 402-579 | 580-1363 |  | This study |
| *Capellomyces foraminis* isolate BGB11 Clone C4 | MT085699 |  |  | 1-220 | 221-401 | 402-578 | 579-1363 |  | This study |
| *Capellomyces elongatus* | MT085701 |  |  | 1-250 | 251-432 | 433-609 | 610-1393 |  | This study |
| *Cyllamyces* |  | 1 (clones 5) | 704 |  |  |  |  | MN1, SP8 |  |
| *Cyllamyces* sp. isolate TSB2 Clone B10 | MT085707 |  |  | 1-170 | 171-347 | 348-537 | 538-1320 |  | This study |
| *Cyllamyces* sp. isolate TSB2 Clone B11 | MT085705 |  |  | 1-170 | 171-347 | 348-538 | 539-1321 |  | This study |
| *Cyllamyces* sp. isolate TSB2 Clone B12 | MT085703 |  |  | 1-170 | 171-347 | 348-537 | 538-1320 |  | This study |
| *Cyllamyces* sp. isolate TSB2 Clone B8 | MT085704 |  |  | 1-168 | 169-344 | 345-536 | 537-1319 |  | This study |
| *Cyllamyces* sp. isolate TSB2 Clone B9 | MT085706 |  |  | 1-168 | 169-345 | 346-535 | 536-1318 |  | This study |
| *Ghazallomyces* |  | 1 (4 clones) | 102 |  |  |  |  |  |  |
| *Ghazallomyces constrictus* isolate AXS31 Clone B1 | MT085693 |  |  | 1-189 | 190-370 | 371-564 | 565-1348 |  | This study |
| *Ghazallomyces constrictus* isolate AXS31 Clone B2 | MT085695 |  |  | 1-186 | 187-364 | 365-556 | 557-1339 |  | This study |
| *Ghazallomyces constrictus* isolate AXS31 Clone B3 | MT085694 |  |  | 1-186 | 187-364 | 365-556 | 557-1339 |  | This study |
| *Ghazallomyces constrictus* isolate AXS31 Clone B5 | MT085696 |  |  | 1-189 | 190-367 | 368-552 | 553-1335 |  | This study |
| *Joblinomyces* |  | 2 (10 clones) | 1076 |  |  |  |  | AL5 |  |
| *Joblinomyces apicalis* isolate GFH681 Clone1 | MT085665 |  |  | 1-213 | 214-388 | 389-561 | 562-1344 |  | This study |
| *Joblinomyces apicalis* isolate GFH681 Clone2 | MT085666 |  |  | 1-213 | 214-390 | 391-564 | 565-1347 |  | This study |
| *Joblinomyces apicalis* isolate GFH681 Clone4 | MT085667 |  |  | 1-215 | 216-393 | 394-568 | 569-1351 |  | This study |
| *Joblinomyces apicalis* isolate GFH681 Clone5 | MT085668 |  |  | 1-215 | 216-393 | 394-568 | 569-1351 |  | This study |
| *Joblinomyces apicalis* isolate GFH681 Clone6 | MT085669 |  |  | 1-213 | 214-389 | 390-563 | 564-1346 |  | This study |
| *Joblinomyces apicalis* isolate GFH683 Clone1 | MT085670 |  |  | 1-213 | 214-389 | 390-563 | 564-1346 |  | This study |
| *Joblinomyces apicalis* isolate GFH683 Clone2 | MT085671 |  |  | 1-213 | 214-389 | 390-563 | 564-1346 |  | This study |
| *Joblinomyces apicalis* isolate GFH683 Clone3 | MT085672 |  |  | 1-212 | 213-388 | 389-562 | 563-1345 |  | This study |
| *Joblinomyces apicalis* isolate GFH683 Clone4 | MT085673 |  |  | 1-213 | 214-389 | 390-563 | 564-1346 |  | This study |
| *Joblinomyces apicalis* isolate GFH683 Clone5 | MT085674 |  |  | 1-213 | 214-389 | 390-563 | 564-1346 |  | This study |
| *Feramyces* |  | 4 (11 clones) | 2373 |  |  |  |  | AL6 |  |
| *Feramyces austinii* isolate DS10 Clone 11 | MG584196.1 |  |  | 1-192 | 193-368 | 369-570 | 571-1353 |  | 28 |
| *Feramyces austinii* isolate DS10 Clone 12 | MG584194.1 |  |  | 1-192 | 193-368 | 369-570 | 571-1353 |  | 28 |
| *Feramyces austinii* isolate DS10 Clone 7 | MG584192.1 |  |  | 1-192 | 193-368 | 369-571 | 572-1354 |  | 28 |
| *Feramyces austinii* isolate DS10 Clone 8 | MG584200.1 |  |  | 1-192 | 193-368 | 369-570 | 571-1352 |  | 28 |
| *Feramyces austinii* isolate DS10 Clone 9 | MG584197.1 |  |  | 1-192 | 193-368 | 369-570 | 571-1353 |  | 28 |
| *Feramyces austinii* isolate F3A Clone 3 | MG584193.1 |  |  | 1-192 | 193-368 | 369-570 | 571-1353 |  | 28 |
| *Feramyces austinii* isolate F3B Clone 10 | MG584190.1 |  |  | 1-192 | 193-368 | 369-570 | 571-1352 |  | 28 |
| *Feramyces austinii* isolate R4A Clone 1 | MG584191.1 |  |  | 1-192 | 193-368 | 369-570 | 571-1353 |  | 28 |
| *Feramyces austinii* isolate R4A Clone 2 | MG584199.1 |  |  | 1-192 | 193-368 | 369-570 | 571-1353 |  | 28 |
| *Feramyces austinii* isolate R4A Clone 3 | MG584198.1 |  |  | 1-192 | 193-368 | 369-570 | 571-1353 |  | 28 |
| *Feramyces austinii* isolate R4A Clone 5 | MG584195.1 |  |  | 1-192 | 193-368 | 369-570 | 571-1353 |  | 28 |
| *Khyollomyces* |  | 1 (1 clone) | 2553 |  |  |  |  | AL1 |  |
| *Khyollomyces ramosus* isolate ZS33 Clone 8 | MT085710 |  |  | 1-193 | 194-369 | 370-543 | 544-1327 |  | This study |
| *Liebetanzomyces* |  | 1 (7 clones) | 31 |  |  |  |  | SP4 |  |
| *Liebetanzomyces* sp. isolate Cel1A Clone 2 | MT085726 |  |  | 1-225 | 226-403 | 404-577 | 578-1361 |  | This study |
| *Liebetanzomyces* sp. isolate Cel1A Clone 3 | MT085727 |  |  | 1-225 | 226-403 | 404-577 | 578-1361 |  | This study |
| *Liebetanzomyces* sp. isolate Cel1A Clone 4 | MT085728 |  |  | 1-225 | 226-403 | 404-577 | 578-1361 |  | This study |
| *Liebetanzomyces* sp. isolate Cel1A Clone 6 | MT085729 |  |  | 1-225 | 226-403 | 404-577 | 578-1362 |  | This study |
| *Liebetanzomyces* sp. isolate Cel1A Clone 7 | MT085730 |  |  | 1-223 | 224-401 | 402-576 | 577-1362 |  | This study |
| *Liebetanzomyces* sp. isolate Cel1A Clone 8 | MT085731 |  |  | 1-225 | 226-403 | 404-577 | 578-1361 |  | This study |
| *Liebetanzomyces* sp. isolate Cel1A Clone 9 | MT085732 |  |  | 1-223 | 224-401 | 402-575 | 576-1359 |  | This study |
| *Neocallimastix* |  | 14 (14 clones) | 794 |  |  |  |  | SP6 |  |
| *Neocallimastix californiae* strain G1** | MCOG01000947.1 |  |  | 12566-12742 | 12743-12917 | 12918-13113 | 13114-13895 |  | 36 |
| *Neocallimastix californiae* strain G1** | MCOG01000947.1 |  |  | 2806-2982 | 2983-3157 | 3158-3354 | 3355-4136 |  | 36 |
| *Neocallimastix californiae* strain G1** | MCOG01001752.1 |  |  | 2193-2369 | 2018-2192 | 1821-2017 | 1820-1036 |  | 36 |
| *Neocallimastix* cf. *cameroonii* isolate G3 | MT085722 |  |  | 1-178 | 179-356 | 357-554 | 555-1338 |  | This study |
| *Neocallimastix* sp isolate HeF5 Clone 1 | MT085723 |  |  | 1-229 | 230-408 | 409-588 | 589-1371 |  | This study |
| *Neocallimastix* sp isolate HeF6 Clone 6 | MT085724 |  |  | 1-240 | 241-419 | 420-602 | 603-1385 |  | This study |
| *Neocallimastix* sp isolate HeF7 Clone 3 | MT085725 |  |  | 1-229 | 230-407 | 408-585 | 586-1368 |  | This study |
| *Neocallimastix* cf. *frontalis* isolate NYF1 | JQ782542.1 |  |  | 67-308 | 309-486 | 487-665 | 666-1390 |  | 23 |
| *Neocallimastix* cf. *frontalis* isolate NYF2 | JQ782543.1 |  |  | 67-307 | 308-485 | 486-672 | 673-1397 |  | 23 |
| *Neocallimastix* cf. *frontalis* isolate NYF3 | JQ782544.1 |  |  | 67-296 | 297-474 | 475-654 | 655-1379 |  | 23 |
| *Neocallimastix* cf. *frontalis* isolate NYF4 | JQ782545.1 |  |  | 67-310 | 311-488 | 489-676 | 677-1401 |  | 23 |
| *Neocallimastix* cf. *frontalis* isolate NYR1 | JQ782546.1 |  |  | 67-307 | 308-486 | 487-669 | 670-1394 |  | 23 |
| *Neocallimastix* cf. *frontalis* isolate NYR2 | JQ782547.1 |  |  | 67-317 | 318-496 | 497-676 | 677-1401 |  | 23 |
| *Neocallimastix* cf. *frontalis* isolate NYR3 | JQ782548.1 |  |  | 67-317 | 318-495 | 496-678 | 679-1403 |  | 23 |
| *Neocallimastix* cf. *frontalis* isolate NYR4 | JQ782549.1 |  |  | 67-295 | 296-473 | 474-654 | 655-1380 |  | 23 |
| *Neocallimastix* cf. *frontalis* isolate NYR5 | JQ782550.1 |  |  | 67-309 | 310-488 | 489-667 | 668-1392 |  | 23 |
| *Oontomyces* |  | 0 | 0 |  |  |  |  |  |  |
| *Oontomyces anksri* strain SSD-CIB1 | JX017310.1 |  |  | 60-291 | 292-467 | 468-642 | 643-695 |  | 26 |
| *Oontomyces anksri* strain SSD-CIB1 | JX017314.1 |  |  |  |  |  | 1-772 |  | 26 |
| *Orpinomyces* |  | 23 clones | 349 |  |  |  |  |  |  |
| *Orpinomyces* sp. OUS1 | AJ864475.1 |  |  | 842-1056 | 1057-1241 | 1242-1418 | 1419-2201 |  | 49 |
| *Orpinomyces* cf. *joyonii* isolate D3A Clone 3 | MT085735 |  |  | 1-183 | 184-360 | 361-539 | 540-1322 |  | This study |
| *Orpinomyces* cf. *joyonii* isolate D3A Clone F11 | MT085736 |  |  | 1-182 | 183-357 | 358-537 | 538-1320 |  | This study |
| *Orpinomyces* cf. *joyonii* isolate D3A Clone G09 | MT085733 |  |  | 1-184 | 185-359 | 360-538 | 539-1321 |  | This study |
| *Orpinomyces* cf. *joyonii* isolate D3A Clone H09 | MT085734 |  |  | 1-187 | 188-362 | 363-540 | 541-1323 |  | This study |
| *Orpinomyces* sp. OYF | JQ782551.1 |  |  | 67-267 | 268-447 | 448-630 | 631-1356 |  | 23 |
| *Orpinomyces* sp. OYR2 | JQ782553.1 |  |  | 65-253 | 254-431 | 432-610 | 611-1335 |  | 23 |
| *Pecoramyces* |  | 2 (4 clones) | 248 |  |  |  |  |  |  |
| *Pecoramyces ruminatium* isolate C1A** | ASRE01020932.1 |  |  | 909-1095 | 1096-1271 | 1272-1452 | 1453-2235 |  | 43 |
| *Pecoramyces ruminatium* isolate C1A** | ASRE01007038.1 |  |  | 790-976 | 977-1152 | 1153-1333 | 1334-2116 |  | 43 |
| *Pecoramyces ruminatium* isolate C1A** | ASRE01022884.1 |  |  | 2760-2946 | 2584-2759 | 2403-2583 | 1620-2402 |  | 43 |
| *Pecoramyces ruminatium* isolate S4B | MT085711 |  |  | 1-184 | 185-360 | 361-542 | 543-1325 |  | This study |
| *Piromyces* |  | 5 (26 clones) | 3818 |  |  |  |  | AL7, UC1 |  |
| *Piromyces finnis* strain finn*** | MCFH01000027.1 |  |  | 1034-1105 | 1106-1285 | 1286-1470 | 1471-2253 |  | 36 |
| *Piromyces finnis* strain finn*** | MCFH01000027.1 |  |  | 9568-9639 | 9640-9819 | 9820-10004 | 10005-10787 |  | 36 |
| *Piromyces finnis* strain finn*** | MCFH01000027.1 |  |  | 18102-18173 | 18174-18353 | 18354-18538 | 18539-19321 |  | 36 |
| *Piromyces* sp. isolate A1 Clone A1 | MT085682 |  |  | 1-199 | 200-375 | 376-548 | 549-1333 |  | This study |
| *Piromyces* sp. isolate A1 Clone A12 | MT085684 |  |  | 1-198 | 199-374 | 375-547 | 548-1330 |  | This study |
| *Piromyces* sp. isolate A1 Clone A2 | MT085679 |  |  | 1-199 | 200-375 | 376-548 | 549-1331 |  | This study |
| *Piromyces* sp. isolate A1 Clone A3 | MT085683 |  |  | 1-199 | 200-375 | 376-548 | 549-1331 |  | This study |
| *Piromyces* sp. isolate A1 Clone A4 | MT085685 |  |  | 1-199 | 200-375 | 376-548 | 549-1331 |  | This study |
| *Piromyces* sp. isolate A1 Clone A5 | MT085688 |  |  | 1-199 | 200-375 | 376-548 | 549-1335 |  | This study |
| *Piromyces* sp. isolate A1 Clone A6 | MT085687 |  |  | 1-199 | 200-375 | 376-548 | 549-1331 |  | This study |
| *Piromyces* sp. isolate A1 Clone A7 | MT085686 |  |  | 1-199 | 200-375 | 376-548 | 549-1334 |  | This study |
| *Piromyces* sp. isolate A1 Clone A8 | MT085681 |  |  | 1-199 | 200-375 | 376-548 | 549-1331 |  | This study |
| *Piromyces* sp. isolate A1 Clone A9 | MT085680 |  |  | 1-199 | 200-375 | 376-548 | 549-1331 |  | This study |
| *Piromyces* sp. isolate Cel1B Clone 1 | MT085717 |  |  | 1-198 | 199-375 | 376-548 | 549-1331 |  | This study |
| *Piromyces* sp. isolate Cel1B Clone 10 | MT085721 |  |  | 1-198 | 199-374 | 375-547 | 548-1332 |  | This study |
| *Piromyces* sp. isolate Cel1B Clone 2 | MT085718 |  |  | 1-198 | 199-375 | 376-548 | 549-1331 |  | This study |
| *Piromyces* sp. isolate Cel1B Clone 3 | MT085719 |  |  | 1-198 | 199-375 | 376-549 | 550-1334 |  | This study |
| *Piromyces* sp. isolate Cel1B Clone 6 | MT085720 |  |  | 1-198 | 199-374 | 375-547 | 548-1332 |  | This study |
| *Piromyces* sp. isolate DB3 Clone B2 | MT085690 |  |  | 1-226 | 227-404 | 405-592 | 593-1350 |  | This study |
| *Piromyces* sp. isolate DB3 Clone B3 | MT085691 |  |  | 1-227 | 228-405 | 406-592 | 593-1359 |  | This study |
| *Piromyces* sp. isolate DB3 Clone B4 | MT085689 |  |  | 1-225 | 226-403 | 404-588 | 589-1351 |  | This study |
| *Piromyces* sp. isolate Jen1 Clone 1 | MT085712 |  |  | 1-201 | 202-378 | 379-552 | 553-1335 |  | This study |
| *Piromyces* sp. isolate Jen1 Clone 2 | MT085713 |  |  | 1-201 | 202-378 | 379-552 | 553-1335 |  | This study |
| *Piromyces* sp. isolate Jen1 Clone 3 | MT085714 |  |  | 1-201 | 202-377 | 378-550 | 551-1333 |  | This study |
| *Piromyces* sp. isolate Jen1 Clone 4 | MT085715 |  |  | 1-201 | 202-377 | 378-550 | 551-1333 |  | This study |
| *Piromyces* sp. isolate Jen1 Clone 5 | MT085716 |  |  | 1-201 | 202-378 | 379-552 | 553-1335 |  | This study |
| *Tahromyces* |  | 4 (4 clones) |  |  |  |  |  |  |  |
| *Tahromyces munnarensis* isolate TDFKJa1924 | MT085677 |  |  | 1-178 | 179-358 | 359-537 | 538-1316 |  | This study |
| *Tahromyces munnarensis* isolate TDFKJa1926 | MT085676 |  |  | 1-178 | 179-358 | 359-537 | 538-1307 |  | This study |
| *Tahromyces munnarensis* isolate TDFKJa1927 | MT085678 |  |  | 1-178 | 179-358 | 359-537 | 538-1320 |  | This study |
| *Tahromyces munnarensis* isolate TDFKJa193 | MT085675 |  |  | 1-178 | 179-358 | 359-537 | 538-1313 |  | This study |
| **Uncultured lineages** |  |  |  |  |  |  |  |  |  |
| **Identified in this study (accession number is the sequence name in Supplementary datasets 1-3)** |  |  |  |  |  |  |  |  |  |
| AL3 | DwGoat_61688 |  | 86 | 57-200 | 201-352 | 353-604 | 605-1358 |  | This study |
| AL4 | Sheep_129918 |  | 1 | 67-268 | 269-425 | 426-630 | 631-1394 |  | This study |
| AL8 | Cow_130070 |  | 151 | 88-273 | 274-432 | 433-653 | 654-1424 |  | This study |
| MN3 | Cow_90808 |  | 3 | 76-280 | 281-438 | 439-640 | 641-1423 |  | This study |
| MN4 | OSUGoat_119881 |  | 3 | 69-276 | 277-434 | 435-666 | 667-1444 |  | This study |
| SK3 | Aoud18_104764 |  | 3 | 67-279 | 280-436 | 437-676 | 677-1444 |  | This study |
| SK4 | Aoud18_141177 |  | 1387 | 71-264 | 265-421 | 422-644 | 645-1421 |  | This study |
| **Not identified in this study (GenBank accession number of a representative ITS1 sequence)** |  |  |  |  |  |  |  |  |  |
| AL2 | GQ826457 |  |  |  |  |  |  |  | 3 |
| BlackRhino | JF423850 |  |  |  |  |  |  |  | 5 |
| DA1 | JX184822 |  |  |  |  |  |  |  | 13 |
| DT1 | GQ850291 |  |  |  |  |  |  |  | 48 |
| JH1/ SP5 | GU911240 |  |  |  |  |  |  |  | 17, 46 |
| KF1 | GQ850345 |  |  |  |  |  |  |  | 45 |
| MN2 | AM690075 |  |  |  |  |  |  |  | 47 |
| SK1 | JF423570 |  |  |  |  |  |  |  | 5 |
| SP1 | GQ678747 |  |  |  |  |  |  |  | 17 |
| SP2 | GQ698377 |  |  |  |  |  |  |  | 17 |
| SP3 | GQ657498 |  |  |  |  |  |  |  | 17 |
| SP7 | GU910219.1 |  |  |  |  |  |  |  | 17 |
| **Novel lineages (accession number is the sequence name in Supplementary datasets 1-3)** |  |  |  |  |  |  |  |  |  |
| RH1 | Cow_130696 |  | 13 | 66-222 | 223-378 | 379-593 | 594-1369 |  | This study |
| RH2 | Oryx_79099 |  | 74 | 67-270 | 271-425 | 426-660 | 661-1440 |  | This study |
| RH3 | AmBis_130671 |  | 3 | 71-246 | 247-404 | 405-613 | 614-1374 |  | This study |
| RH4 | Cow_156860 |  | 13 | 68-238 | 239-393 | 394-597 | 598-1379 |  | This study |
| RH5 | Sheep_119174 |  | 1681 | 69-280 | 281-438 | 439-646 | 647-1426 |  | This study |
| RH6 | Cow_144271 |  | 2 | 78-289 | 290-446 | 447-653 | 654-1433 |  | This study |

\*GenBank Accession numbers are provided for Sanger sequenced clones from all fungal isolates. PacBio-generated datasets are present in GenBank in the Bioproject accession number PRJNA609702, Biosample accession numbers SAMN14258225, and Targeted Locus Study project accession KDVX00000000. FASTA files of ITS1-5.8S-ITS2-D1/D2 LSU region, as well as bioinformatically extracted ITS1 region and D1/D2 LSU regions are provided as supplementary documents (Datasets 1-3)
\*\* Sequences extracted from a genomic assembly

### D1/D2 LSU versus ITS1 as a taxonomic marker

#### Intra-genus length variability

The ITS1 and D1/D2 LSU regions were bioinformatically extracted from the 116 Sanger-generated clone sequences from this study and previous studies^23, 28, 29, 49^ (accession numbers in Table 1), the rRNA loci from two Neocallimastigomycota genomes (*Pecoramyces ruminantium* strain C1A, and *Neocallimastix californiae* strain G1)^36, 43^ in which the entire rrn operon sequence data is available, and from the PacBio-generated environmental amplicons in this study. The ITS1 region displayed a high level of length heterogeneity, ranging in size between 141 and 250 bp (median 191 bp, Figure 1a), with 75% of sequences ranging between 182-208 bp in length. Some genera had shorter than median ITS1 region length, e.g. *Cyllamyces* (range 141-173 bp), *Buwchfawromyces* (range 155-169 bp), and candidate genus AL3 (range 145-148 bp), while others exhibited a longer than median ITS1 region length, e.g. *Liebetanzomyces* (range 198-225 bp), and candidate genus RH5 (range 191-224 bp) (Figure 1a). A third group of genera displayed a wide range of length heterogeneity, e.g. *Neocallimastix* (range 160-244 bp), *Caecomyces* (range 192-250), and *Piromyces* (range 173-225 bp). Few genera and candidate genera displayed a fairly narrow range of ITS1 length, e.g. AL3 (141-148 bp), but this is potentially a reflection of the paucity of sequences belonging to these genera obtained in this study (Figure 1a).

**Figure 1.**
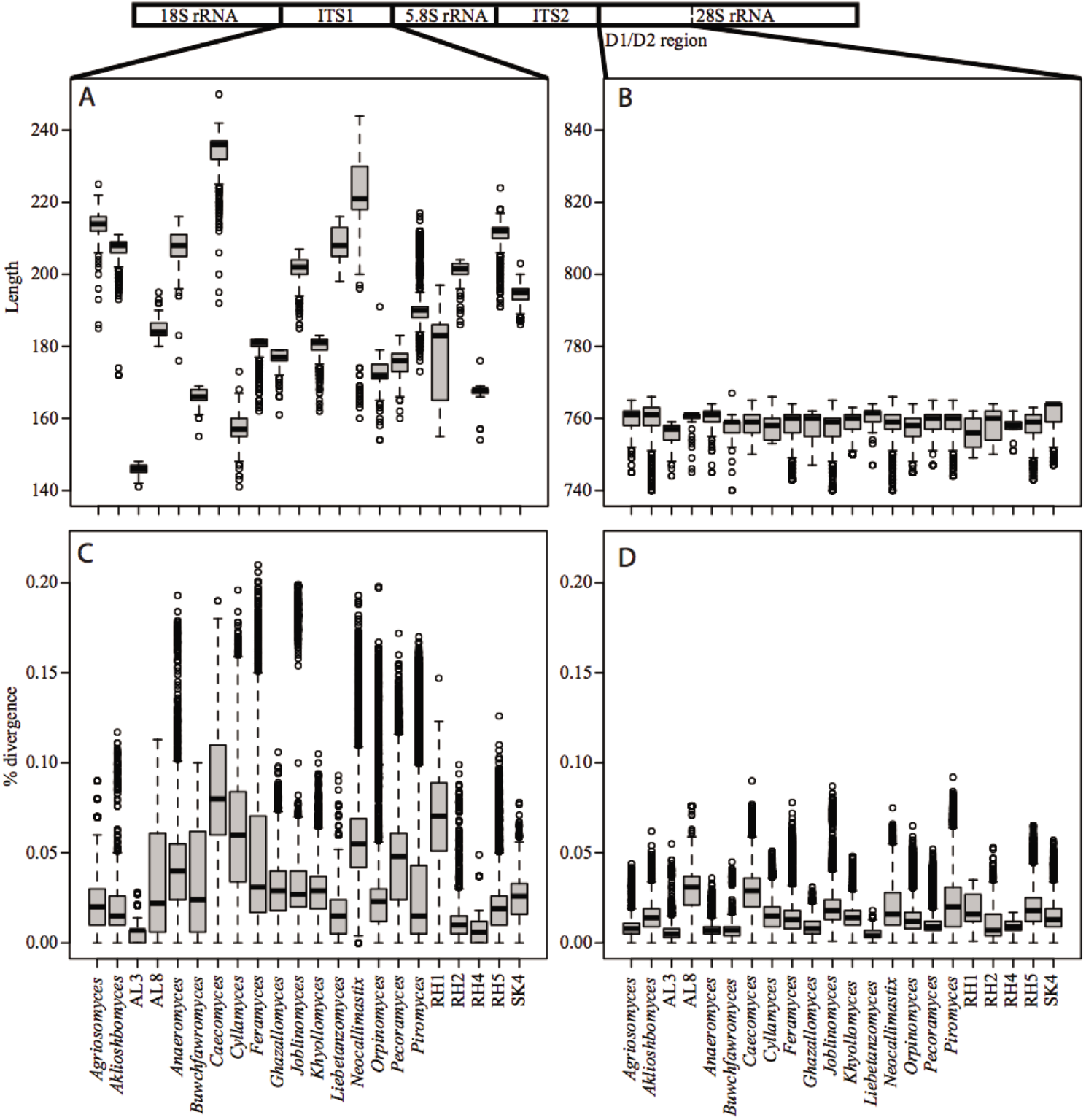
Box and whisker plots showing the variability in intra-genus length (A-B) and sequence divergence cutoff (C-D) for the ITS1 (A, C) and D1/D2 LSU (B, D) regions. A cartoon of the rRNA locus is shown on top. Genera and candidate genera with at least 5 sequences encompassing the full region “ITS1-5.8S-ITS2-D1/D2 LSU” were used to construct this plot as detailed in the methods section. The candidate genera AL4, MN3, MN4, RH3, RH6, and SK3 had only a few sequence representatives (1-3) and so are not included in the plot.

On the other hand, a much lower level of length heterogeneity was identified in the D1/D2 LSU (Figure 1b), ranging in size between 740-767 bp (median 760 bp), and where 75% of the sequences ranged between 757-761 bp, with all genera consistently displaying a much narrower D1/D2-LSU length heterogeneity, ranging between 11 bp in RH4 and 26 bp in the genera *Neocallimastix* and *Aklioshbomyces*.

#### Intra-genus sequence divergence

The ITS1 region displayed intra-genus sequence divergence ranging from 0.4 to 21% (median 3.2%), with 75% of the pairwise divergence values ranging between 1.7-6%. Genera displaying the highest level of divergence were *Caecomyces* (1-18.9%, median 8.3%), *Cyllamyces* (0.6-19.6%, median 5.5%), and *Neocallimastix* (0.4-19.3%, median 5.5%) (Figure 1c). On the other hand, intra-genus sequence divergence of the D1/D2 LSU ranged between 0.1-9.2% (median 1.4%), with 75% of the pairwise divergence values ranging between 0.8-2.1%. Genera displaying highest level of divergence were *Feramyces* (0.1-7.8%), *Joblinomyces* (0.1-8.7%, *Caecomyces* (0.1-9%), and *Piromyces* (0.1-9.2%) (Figure 1d).

#### Within strain length variability

Within strain length heterogenicity examined in 19 strains with 2 or more sequenced clones ranged between 0-5 bp (Figure 2a) for ITS1 region and 0-1 bp for the LSU region (Figure 2b).

**Figure 2.**
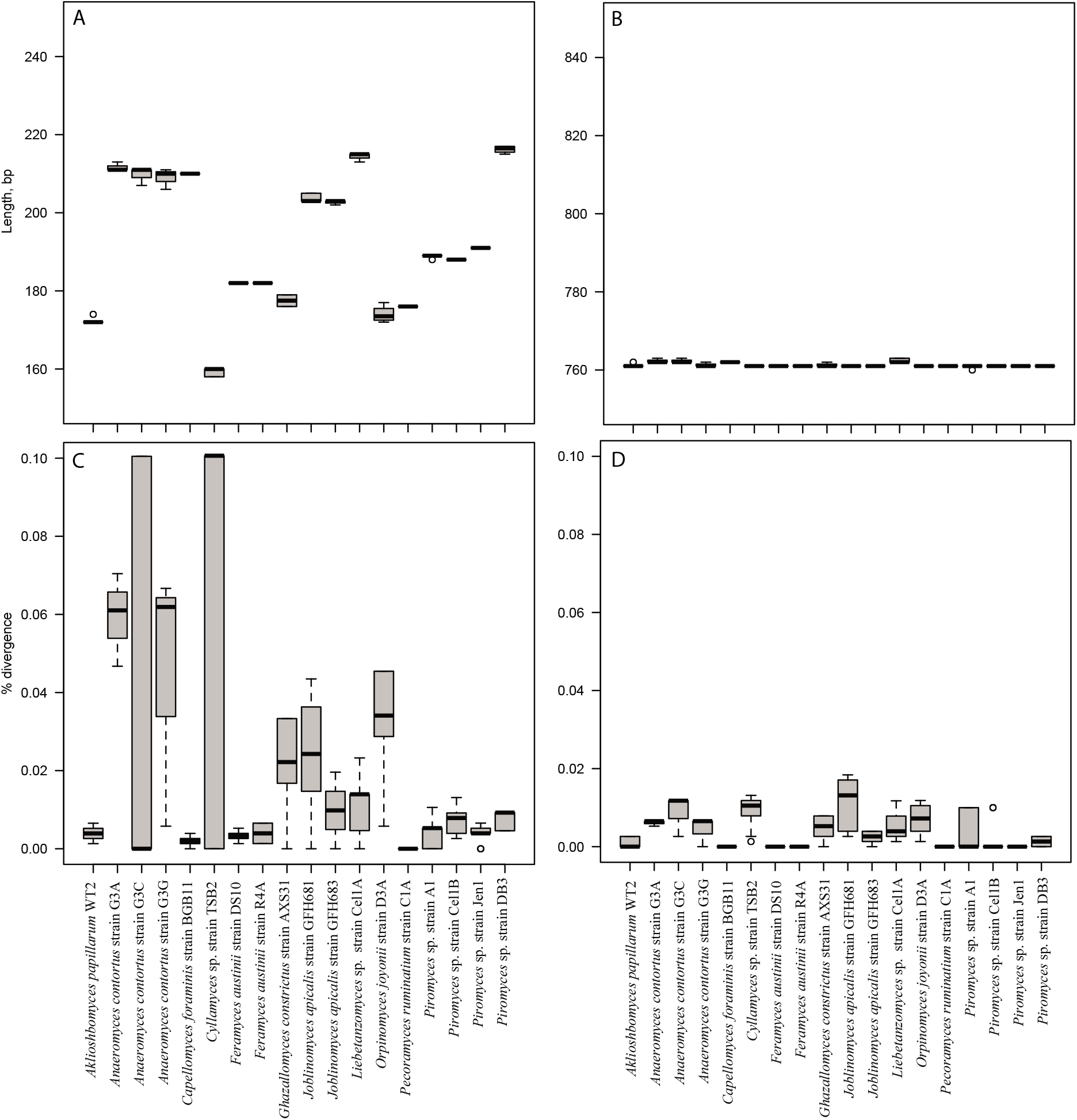
Box and whisker plots showing the variability in intra-strain length (A-B) and sequence divergence cutoff (C-D) for the ITS1 (A, C) and D1/D2 LSU (B, D) regions.

#### Within strain sequence divergence

Examining the 19 strains with more than two sequenced clones, the full ITS1 region showed intra-strain sequence divergence ranging from 0.1-10.01% (Figure 2c). Similar, and even higher levels of intra-strain ITS1 variability was previously reported e.g. up to 12.9% in *Buwchfawromyces eastonii* strain GE09^15^. On the other hand, within strain D1/D2 LSU rRNA region showed a much lower sequence divergence, ranging from 0.13-1.84% (Figure 2d).

### Neocallimastigomycota diversity assessment using D1/D2 LSU as a phylogenetic marker

#### Phylogenetic diversity and Novelty

A total of 17,697 high-quality long-read amplicons were obtained. Phylogenetic analysis using the D1/D2 LSU amplicons assigned all sequences into 28 different genera/candidate genera (Figure 3a, Figure 4a, Figure S1) and 298 species level OTUs_0.02_. AGF genera identified in this study included members of the previously described genera *Anaeromyces, Buwchfawromyces, Caecomyces*, *Cyllamyces*, *Liebetanzomyces*, *Neocallimastix*, *Orpinomyces*, *Pecoramyces*, and *Piromyces*. In addition, sequences representing multiple novel genera were also identified (Figure 3a, Figure 4a, Figure S1), some of which have been subsequently isolated, named, and characterized in separate publications, e.g. *Feramyces*^28^, *Aklioshbomyces*, *Agriosomyces*, *Ghazallomyces*, and *Khyollomyces*^50^. Finally, six novel candidate genera were identified and designated RH1-RH6 (Figure 3a, Figure 4a, Figure S1). All of these six novel genera were encountered in extremely low abundance in a few samples (Figure 3a), with the notable exception of RH5, which was present in high relative abundance in multiple animals e.g. domesticated sheep (96.22%), blackbuck deer (52.41%), axis deer (20.71%), and an aoudad sheep sample (11.75%).

**Figure 3.**
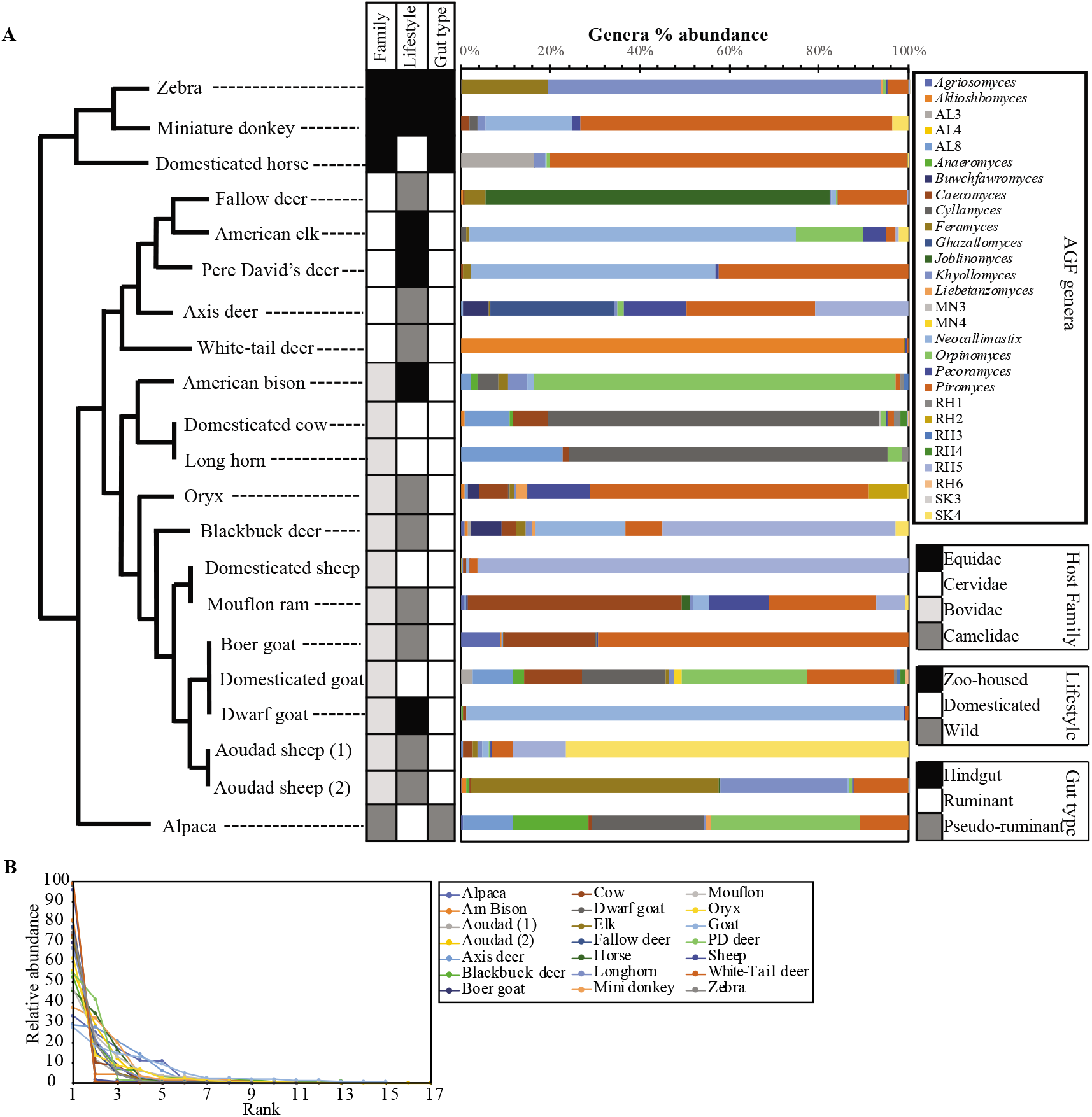
AGF genera distribution across the animal studied. (A) Percentage abundance of AGF genera in the animals studied. The tree is intended to show the relationship between the animals and is not drawn to scale. Host phylogeny (family), lifestyle, and gut type are shown for each animal. The X-axis shows the percentage abundance of AGF genera. (B) Rank abundance curves are displayed for each animal showing a distribution pattern in which a few genera (1-5) represent the majority (>10%) of the sequences obtained.

**Figure 4.**
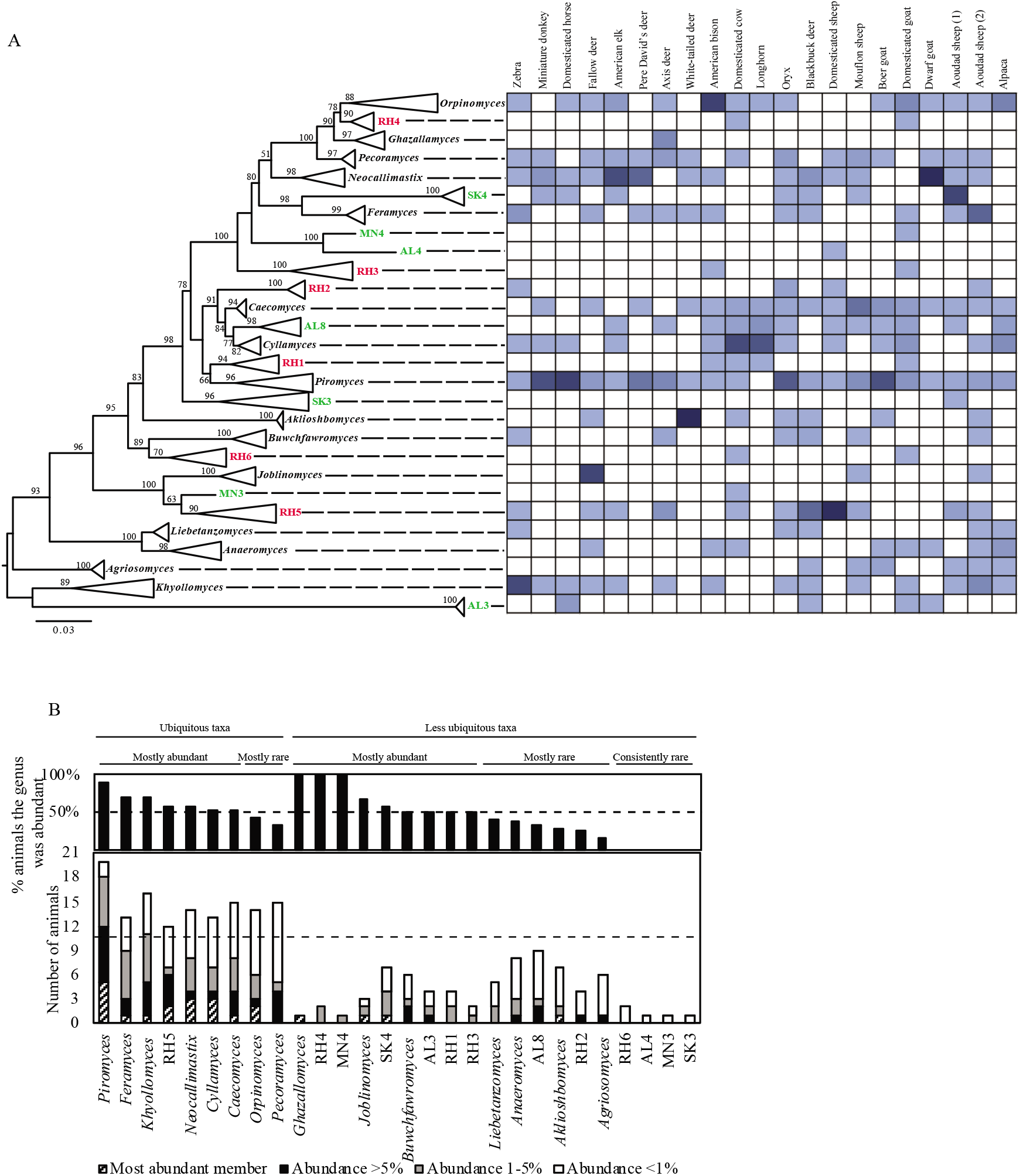
(A) Phylogenetic tree constructed using the D1/D2 LSU sequences of representatives of each of the 28 genera/candidate genera identified in this study. Sequences were aligned using the MAFFT aligner and maximum likelihood tree was constructed in FastTree^61, 62^. Bootstrap values are based on 100 replicates and are shown for branches with >50% bootstrap support. Genera with cultured representatives are shown in black, uncultured candidate genera identified in previous ITS1-based studies are shown in green, while the 6 novel genera identified in the current study are shown in red. The distribution of each of these genera/candidate genera in the animals studied is shown as a heatmap on the right. (B) AGF genera distribution patterns. The total number of animals harboring each of the genera identified in this study is shown on the Y-axis, with the different colored stacked bars reflecting the number of animals where the genus was the most abundant member, occurred with high (>5%) abundance, occurred with medium (1-5%) abundance, or occurred with low (<1%) abundance. AGF genera are classified into one of the five distribution patterns shown on top of the graph using empirical cutoffs for ubiquity (presence in at least 50% of the animals studied, shown as the dotted line across the bottom bar chart), as well as the fraction of animals where the genus abundance was above 1% (shown as the top bar chart).

#### Diversity estimates, and distribution patterns

The number of AGF genera encountered per sample varied widely from 5 (in Pere David’s deer, and Longhorn cattle) to 16 (in one Aoudad sheep sample) (Table 2, Figure 3a, Figure 4a). However, in each of these samples a distribution pattern was observed in which a few genera represent the absolute majority of the sequences obtained. Excluding genera present in less than 1% abundance would lower the number of genera encountered per animal to 1 (in white-tail deer and dwarf goat) −10 (domesticated goat). Usually, 1-5 taxa were present in >10% abundance per animal (Figure 3b).

**Table 2.** Animals sampled in this study, numbers of sequences obtained (N), number of observed OTUs, and various diversity indices both at the species equivalent (0.02) and the genus levels

| Sample | Host description |  | N* | Observed number of OTUs |  | Chao |  | Ace |  | Simpson evenness |  | Shannon |  | Diversity ranking** |  | Coverage*** |  |
| --- | --- | --- | --- | --- | --- | --- | --- | --- | --- | --- | --- | --- | --- | --- | --- | --- | --- |
|  | Family | Lifestyle |  | Sp. Eq. | Genus | Sp. Eq. | Genus | Sp. Eq. | Genus | Sp. Eq. | Genus | Sp. Eq. | Genus | Sp. Eq. | Genus | Sp. Eq. | Genus |
| Alpaca | Camelidae | Domestic | 240 | 19 | 9 | 26.2 | 9.5 | 30.5 | 10.8 | 0.27 | 0.50 | 1.99 | 1.64 | 17.3 | 18.8 | 0.94 | 0.99 |
| American bison | Bovidae | Zoo | 183 | 17 | 11 | 22.6 | 11.3 | 41.6 | 12.2 | 0.13 | 0.14 | 1.45 | 0.90 | 9.8 | 7.5 | 0.93 | 0.99 |
| American elk | Cervidae | Zoo | 99 | 11 | 9 | 16 | 11 | 28.2 | 15.3 | 0.17 | 0.21 | 1.12 | 1.01 | 6.3 | 10.5 | 0.92 | 0.96 |
| Aoudad sheep (1) | Bovidae | Wild | 3381 | 80 | 17 | 111 | 23 | 159.4 | 27 | 0.03 | 0.15 | 1.54 | 1.17 | 13.5 | 7 | 0.99 | 1 |
| Aoudad sheep (2) | Bovidae | Wild | 1779 | 55 | 13 | 57 | 13 | 59.9 | 13.4 | 0.05 | 0.13 | 1.78 | 0.91 | 11.2 | 11.8 | 0.99 | 1 |
| Axis deer | Cervidae | Wild | 367 | 18 | 9 | 18.6 | 9.5 | 19.6 | 11.6 | 0.36 | 0.49 | 2.17 | 1.61 | 19 | 19.5 | 0.99 | 0.99 |
| Blackbuck deer | Bovidae | Wild | 145 | 21 | 13 | 34.8 | 16 | 67.7 | 17 | 0.18 | 0.24 | 1.93 | 1.59 | 16 | 16.7 | 0.91 | 0.97 |
| Boer goat | Bovidae | Wild | 2503 | 41 | 9 | 49.1 | 9 | 58.8 | 9 | 0.05 | 0.21 | 1.12 | 0.88 | 6.5 | 8.3 | 0.99 | 1 |
| Domestic cow | Bovidae | Domestic | 727 | 45 | 13 | 46.7 | 16 | 48.5 | 15.4 | 0.06 | 0.14 | 1.95 | 1.03 | 13.5 | 10.2 | 0.98 | 1 |
| Domestic goat | Bovidae | Domestic | 162 | 23 | 15 | 39.5 | 16 | 70.7 | 17.2 | 0.40 | 0.43 | 2.52 | 2.10 | 20.3 | 21 | 0.87 | 1 |
| Domestic horse | Equidae | Domestic | 498 | 15 | 8 | 15.5 | 8.3 | 17.7 | 10 | 0.11 | 0.35 | 0.94 | 1.18 | 5.3 | 13.7 | 0.99 | 1 |
| Domestic sheep | Bovidae | Domestic | 1349 | 33 | 9 | 33.7 | 9 | 34.5 | 9.5 | 0.04 | 0.12 | 0.72 | 0.25 | 2.5 | 3 | 1 | 1 |
| Dwarf goat | Bovidae | Zoo | 519 | 15 | 8 | 20.3 | 18 | 36.7 | 17.5 | 0.08 | 0.13 | 0.49 | 0.14 | 1 | 2 | 0.98 | 0.99 |
| Fallow deer | Cervidae | Wild | 1368 | 43 | 12 | 46.7 | 12.3 | 48.7 | 13.2 | 0.08 | 0.13 | 1.67 | 0.78 | 14.5 | 5.3 | 0.99 | 1 |
| Longhorn cattle | Bovidae | Domestic | 62 | 9 | 5 | 16.5 | 5 | 47.2 | 5.8 | 0.28 | 0.41 | 1.37 | 0.96 | 11 | 10 | 0.82 | 0.98 |
| Miniature donkey | Equidae | Zoo | 56 | 12 | 8 | 30 | 11 | 191.6 | 18 | 0.23 | 0.46 | 1.50 | 1.46 | 11 | 17.2 | 0.80 | 0.93 |
| Mouflon ram | Bovidae | Wild | 297 | 17 | 11 | 35 | 11.3 | 111 | 14.3 | 0.23 | 0.31 | 1.80 | 1.55 | 16 | 17.2 | 0.95 | 0.99 |
| Oryx | Bovidae | Wild | 780 | 34 | 15 | 36.2 | 16 | 41.9 | 17.7 | 0.26 | 0.16 | 2.52 | 1.35 | 20.7 | 13.8 | 0.98 | 0.98 |
| Pere David's deer | Cervidae | Zoo | 169 | 7 | 6 | 10 | 6.5 | 17 | 7.8 | 0.32 | 0.36 | 0.96 | 0.88 | 6.7 | 10.5 | 0.96 | 0.99 |
| White-tail deer | Cervidae | Wild | 946 | 23 | 6 | 23 | 7.5 | 23.4 | 14.7 | 0.06 | 0.17 | 0.75 | 0.07 | 2.8 | 1 | 1 | 1 |
| Zebra | Equidae | Zoo | 2067 | 55 | 11 | 73 | 13 | 85.4 | 15 | 0.03 | 0.15 | 1.10 | 0.76 | 6 | 6 | 0.99 | 1 |

\*: N refers to the number of sequences obtained for each of the animals sampled.
\*\*: Diversity ranking is the average rank obtained using both an information-related diversity ordering method (Renyi generalized entropy), and an expected number of species-related diversity ordering method (Hulbert family of diversity indices). Samples are ranked from the least diverse (rank 1) to the most diverse (rank 21).
\*\*\*: Coverage refers to the Good's coverage index.

Using empirical cutoffs for ubiquity (presence in at least 50% of the animals studied) and abundance (above 1%), we identify five different distribution patterns for AGF genera encountered in this study (Figure 4b); 1. Ubiquitous mostly abundant genera: These are the genera identified in at least 50% of the animals studied and where their relative abundances exceed 1% in at least 50% of their hosts: This group includes *Piromyces, Feramyces, Khyollomyces*, RH5, *Neocallimastix, Cyllamyces*, and *Caecomyces*. 2. Ubiquitous mostly rare genera: These are the genera identified in at least 50% of the animals studied and where their relative abundances were lower than 1% in at least 50% of their hosts. This group includes *Orpinomyces*, and *Pecoramyces*. 3. Less ubiquitous but mostly abundant genera: These are the genera identified in < 50% of the animals studied but where their relative abundances exceed 1% in at least 50% of their hosts. This group includes *Ghazallomyces*, RH4, MN4, *Joblinomyces*, SK4, *Buwchfawromyces*, AL3, RH1, and RH3. 4. Less ubiquitous mostly rare genera: These are the genera identified in < 50% of the animals and where their relative abundances were lower than 1% in at least 50% of their hosts. This group includes *Liebetanzomyces, Anaeromyces*, AL8, *Aklioshbomyces*, RH2, and *Agriosomyces*. 5. Less ubiquitous consistently rare genera: These are the genera identified in < 50% of the animals and where their relative abundances never exceeded 1% in any of their hosts. This group includes RH6, AL4, MN3, and SK3.

Multiple diversity estimates (number of observed genera, Chao and Ace richness estimates, Shannon diversity index, Simpson evenness, as well as diversity rankings) were computed for each sample (Table 2). The highest genus-level richness was observed in aoudad sheep, dwarf goat, oryx, domesticated cow, domesticated goat, miniature donkey, zebra, and blackbuck deer samples, while the highest genus-level diversity (based on diversity ranking and Shannon index) was observed in domesticated goat, alpaca, axis deer, blackbuck deer, mouflon ram, miniature donkey, oryx, and domesticated horse. On the other hand, the lowest genus-level richness was observed in longhorn cattle, Pere David’s deer, Boer goat, domesticated horse, domesticated sheep, and alpaca, while the lowest genus-level diversity was observed in Fallow deer, zebra, domesticated sheep, dwarf goat, and white-tail deer.

When correlated to animal host phylogeny or animal lifestyle (24 possible combinations), all diversity estimates showed low correlation coefficients (Cramer’s V statistic <0.49) at both the genus and the species equivalent levels (Table S1). Student t-tests were used to examine the significance of the difference in diversity estimates at the genus and species equivalent levels between animal host families (families Bovidae, Cervidae, and Equidae) as well as animal lifestyle (zoo-housed, wild, and domesticated). Only three of these showed a significant difference (Student t-test p-value <0.05): Family Bovidae had a significantly higher observed number of genera and significantly higher Chao estimate at the genus level, and zoo-housed animals had significantly lower Shannon diversity at the species equivalent level (Table S1).

#### Community structure

We used a combination of ordination methods and Student t tests to identify associations between AGF genera and host factors. Non-metric multidimensional scaling based on the genus-level Bray-Curtis indices (Figure 5a-b) identified a few patterns. The genera *Aklioshbomyces, Ghazallomyces, Joblinomyces, Feramyces, Buwchfawromyces*, and *Pecoramyces* seem to be more prevalent in some wild animals (e.g. black buck deer, mouflon, oryx, axis deer, and white tailed deer; filled squares in Figure 5a), while some zoo-housed animals (e.g. elk, dwarf goat, and miniature donkey; grey squares in Figure 5a) clustered together based on the abundance of *Neocallimastix*, *Caecomyces*, and *Liebetanzomyces*. Few domesticated animals (e.g. domesticated goat, longhorn, alpaca, and domesticated cow; open squares in Figure 5a) clustered together based on the abundance of *Cyllamyces*, AL8, MN3, MN4, RH1, RH3, RH4, and RH6. Animal host family had a slightly less apparent effect on AGF community structure (Figure 5b) with the exception of the importance of *Aklioshbomyces* and *Ghazallomyces* in family Cervidae, and AL3 and *Khyollomyces* in family Equidae.

**Figure 5.**
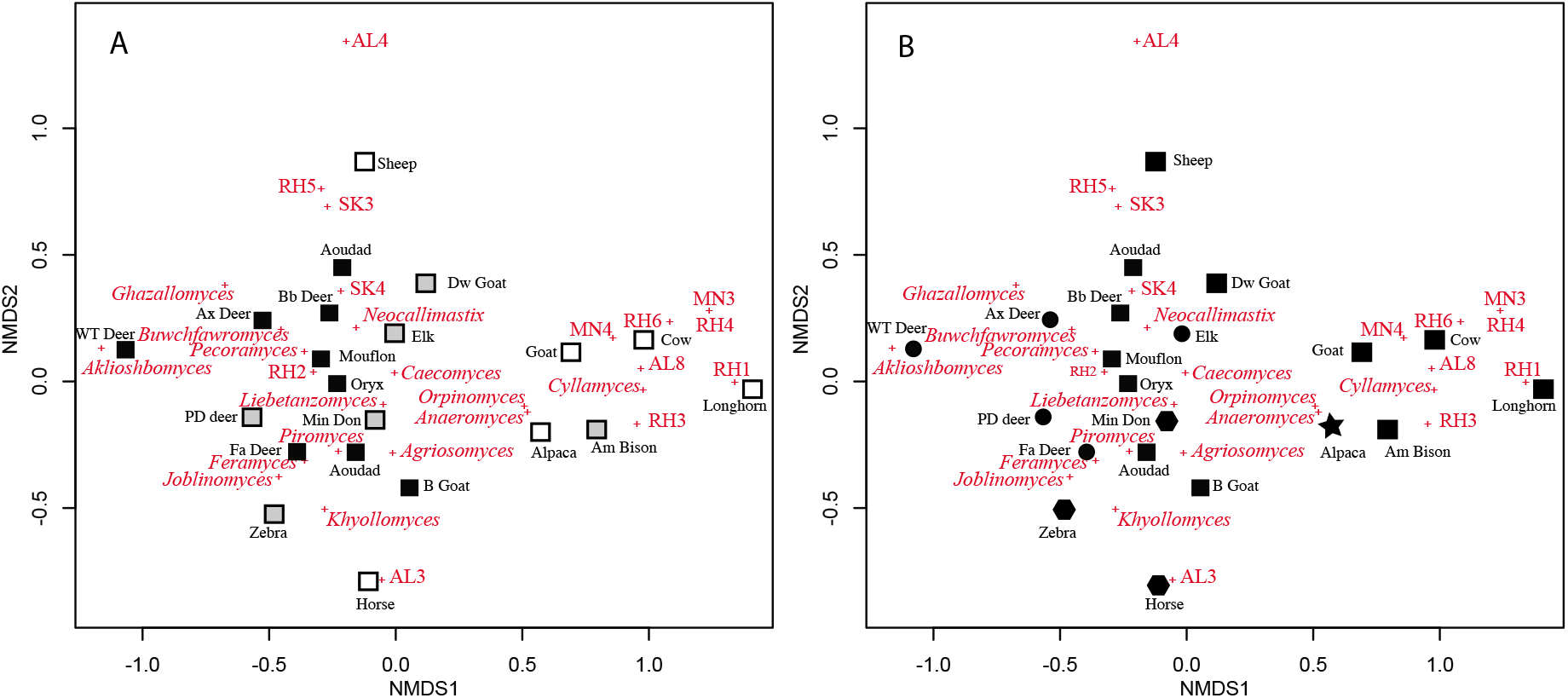
Nonmetric multidimensional scaling based on pairwise Bray-Curtis dissimilarity indices at the genus level. Samples are shown as symbols and displayed in black text while AGF genera are shown as ‘+’ and displayed in red text. (A) Symbols reflect lifestyle with domesticated animals shown as white squares, zoo-housed animals shown as grey squares, and wild animals shown as black squares. (B) Symbols reflect animal host phylogeny with family Bovidae shown as squares, family Cervidae shown as circles, family Equidae shown as hexagons, and family Camelidae shown as a star. Abbreviations: Am Bison, American bison; Ax deer, Axis deer; B goat, Boer goat; Bb deer, Blackbuck deer; Dw Goat, Dwarf goat; Fa deer, Fallow deer; Min Don, Miniature donkey; PD deer, Pere David’s deer; WT deer, White-tail deer.

To test the significance of these observed patterns, Student t-tests were used to identify significant associations between specific AGF taxa and host phylogeny (families Bovidae, Equidae, Cervidae), or animal lifestyle (zoo-housed, domesticated, wild). From all possible associations (168 total; 28 genera x 3 host families and 3 lifestyles), significant differences were observed only in the following cases. The AGF genera AL3, *Khyollomyces*, and *Piromyces* were significantly more abundant in family Equidae (p-value=0.014, 0.018, and 0.034 respectively), while the genera *Aklioshbomyces*, *Ghazallomyces*, and *Joblinomyces* were significantly more abundant in family Cervidae (p-value=0.074, 0.072, and 0.075 respectively). On the other hand, the animal lifestyle had slightly more significant effect on AGF community structure as follows: The genus *Neocallimastix* was significantly more abundant in zoo-housed animals (p-value=0.007), the genera *Aklioshbomyces, Buwchfawromyces*, and *Pecoramyces* were significantly more abundant in wild animals (p-value=0.047, 0.028, and 0.014 respectively), and the genera *Cyllamyces*, AL8, RH1, RH4, and RH6 were significantly more abundant in domesticated animals (p-value=0.001, 0.001, 0.011, 0.018, and 0.054 respectively). Finally, for individual animals species with enough replication in our study, the genera *Cyllamyces*, AL8, and RH1 were significantly more abundant in *Bos taurus* (p-values=1.86E-11, 3E-5, and 2.27E-9, respectively), the genera *Caecomyces* and RH5 were significantly more abundant in *Ovis aries* (p-values=0.006, and 0.004 respectively), and the genera *Feramyces* and SK4 were significantly more abundant in *Ammotragus lervia* (p-values=0.002, and 0.0006, respectively). Further, some genera were only encountered in one animal, demonstrating a probable strong AGF genus-host preference. These genera include *Ghazallomyces* only encountered in axis deer, AL4 only encountered in domesticated sheep, MN3 only encountered in domesticated cow, and MN4 only encountered in domesticated goat.

### Neocallimastigomycota isolation

A total of 216 AGF isolates were obtained from 21 animals (Table 3). Success in isolation and maintenance of that large number of isolates was enabled by the implementation of various techniques for isolation, and the development of a reliable storage procedure^51^. Isolates obtained belonged to 12 different genera (Table 3), six of which were exclusively isolated in this study, and characterized in separate publications (*Akhlioshbomyces*, *Ghazallomyces*, *Capellomyces, Agriosomyces*, *Khoyollomyces* (AL1), and *Feramyces* (AL6)^28, 50^. In general, 1-3 AGF genera were isolated per sample. Isolation efforts captured anywhere between 6.3% (1 of 16 genera) to 27.3% (3 of 11 genera) of AGF genera identified in a single sample using culture-independent D1/D2 LSU gene-based analysis. However, these values are highly affected by the fact that sequencing efforts are capable of identification of AGF genera present in extremely low levels of relative abundance. Indeed, excluding rare taxa (those present at <1% abundance), the culturability goes up to 10% (1 of 10 genera)-100% (2 of 2 genera).

**Table 3.** Number and sources of isolates obtained in thus study.

| AGF genus | Source | Number of isolates | % Abundance in that animal |
| --- | --- | --- | --- |
| *Agriosomyces* | Mouflon | 4 | 3.28 |
|  | Boer goat | 1 | 8.5 |
| *Aklioshbomyces* | White-tail deer | 9 | 98.95 |
| *Anaeromyces* | Domesticated cow | 4 | 0.68 |
|  | domesticated goat | 12 | 2.44 |
|  | American bison | 4 | 1.63 |
|  | Alpaca | 4 | 17.01 |
| *Caecomyces* | Fallow deer | 1 | 0.44 |
| *Capellamyces* | Boer goat | 5 | ND |
| *Feramyces* | Aoudad sheep (1) | 5 | 55.31 |
|  | Fallow deer | 1 | 4.46 |
| *Ghazallomyces* | Axis deer | 11 | 27.79 |
| *Khyollomyces* | Zebra | 16 | 74.5 |
| *Neocallimastix* | Dwarf goat | 7 | 97.88 |
|  | Fallow deer | 10 | 1.24 |
|  | Pere David's deer | 10 | 54.44 |
|  | American elk | 12 | 72 |
| *Orpinomyces* | Domesticated cow | 8 | 1.09 |
|  | Longhorn | 3 | 3.03 |
|  | American bison | 6 | 80.43 |
|  | Alpaca | 6 | 33.2 |
| *Pecoramyces* | Domesticated sheep | 10 | 0.37 |
|  | Mouflon | 3 | 12.79 |
|  | Oryx | 11 | 13.78 |
|  | Aoudad sheep (2) | 13 | 0.39 |
| *Piromyces* | Domesticated cow | 5 | 1.64 |
|  | domesticated sheep | 3 | 1.6 |
|  | Mouflon ram | 2 | 23.61 |
|  | Axis deer | 1 | 28.88 |
|  | Blackbuck deer | 7 | 6.21 |
|  | Domesticated horse | 6 | 80.12 |
|  | Miniature donkey | 16 | 69.64 |

We sought to determine how community structure and isolation efforts correlate, and whether obtaining isolates belonging to a specific genus could be predicted from the community structure of the sample. We observed a strong Pearson correlation (r=0.79) between the abundance of a certain genus in a sample and the frequency of its isolation. On the other hand, the success of isolation of the most dominant member of the community was negatively affected by the sample evenness (Pearson correlation coefficient = −0.87). Indeed, our ability to isolate the novel genera *Aklioshbmyces, Ghazallomycota*, and *Khyollomyces* could be attributed to their presence in high relative abundance in samples from which they were recovered (Table 3), as opposed to their rarity/absence in other samples (Figure 3, 4). Within ubiquitous genera, we observed that the abundance-success of isolation correlation described above is stronger for monocentric taxa (Pearson correlation coefficients= 0.83, 0.96, 0.92, and 1 for *Pecoramyces, Feramyces*, *Neocallimastix*, and *Agriosomyces*, respectively), while such relationship was much weaker in polycentric taxa (Pearson correlation coefficients= 0.31, and 0.58 for *Orpinomyces*, and *Anaeromyces*, respectively). However, the polycentric nature of these genera (ability to propagate even in the absence of zoospore production, and the larger colony size on roll tubes) enabled their isolation even when they constituted a minor fraction of the total community.

## Discussion

### LSU as a phylogenetic marker for AGF diversity surveys

We highlight and quantify the advantages associated with the utilization of D1/D2 LSU as a phylomarker for the AGF when compared to the currently utilized ITS1 region (Figures 1, 2). We also report on overcoming the three main hurdles (lack of reference sequences from uncultured genera, correlating D1/D2 LSU data to currently available ITS1 datasets, and amplicon length precluding utilization of Illumina platform) associated with D1/D2 LSU use as a phylomarker. To address the lack of reference LSU sequence data, we undertook a multi-year isolation effort to provide a comprehensive D1/D2-LSU database from a wide range of AGF taxa, a necessary approach given the lack of LSU sequence data from multiple historic taxa, unavailability of AGF in culture collections, and difficulties in maintenance of this fastidious group of organisms. To correlate D1/D2 LSU data to currently available ITS1 datasets and overcome amplicon length constrains, we utilized a SMRT-PacBio sequencing approach to obtain sequences comprising the region spanning from the start of the ITS1 region to the end of the D1/D2-LSU region in the rRNA locus (1400-1500 bp). In the process, we not only increased the representation of D1/D2-LSU sequences from all cultured taxa, but also identified D1/D2-LSU sequences of yet-uncultured taxa previously defined only by their ITS1 sequences (e.g. AL3, AL4, AL8, MN3, MN4, SK3, and SK4), as well as defined 6 completely novel AGF candidate genera (RH1-RH6). Collectively, this dataset (17,697 sequences of environmental D1/D2 LSU annotated by their taxonomy (Dataset 3), plus 116 Sanger-generated clone sequences and genomic rRNA loci sequences (Table 1 with accession numbers) could pave the way for future D1/D2 LSU-based AGF diversity surveys. We anticipate that additional sampling and culture-independent studies using the whole region, as well as future isolation efforts will identify the corresponding D1/D2-LSU region for those few yet-uncultured ITS1-defined lineages that we failed to capture in our study.

Single molecule real-time (SMRT) PacBio sequencing technology enables long read sequencing by a single uninterrupted DNA polymerase molecule. The SMRT sequencing protocol involves ligating hairpin adaptors to the ends of double-stranded DNA (PCR products in the case of culture-independent studies), leading to the circularization of the DNA. This subsequently allows the sequencing polymerase to pass around the molecule multiple times. The re-sequencing by multiple passages increases sequence coverage thereby significantly reducing error rates from initial values of up to 15%, to levels lower than 1%. Culture-independent studies in bacteria, archaea, and fungi^33, 52, 53, 54^ have successfully applied the technology. We, here, provide the basis for its application to culture-independent studies in anaerobic gut fungi. We applied rigorous control to ensure the high quality of reads utilized to build the single molecule consensus read sequences (by using a minimum threshold of 5 full passes and 99.95% predicted accuracy), followed by pre-processing in Mothur to remove sequences with ambiguities or an average quality score below 25. Also, we anticipate that future AGF diversity studies employing PacBio sequencing of the D1/D2-LSU region (rather than the full ITS1-5.8S-ITS2-D1/D2 LSU region) would be further enabled by the shorter amplicon length (~700 as opposed to ~1300-1400 bp), as well as recent (e.g. Sequel II) and future anticipated improvements in SMRT sequencing technology.

### Discovery and characterization of novel AGF lineages

D1/D2 LSU-based diversity assessment of 21 fecal samples identified multiple novel AGF candidate genera (Figure 3, 4), five of which were subsequently isolated and described in separate publications (*Feramyces*^28^, *Aklioshbomyces*, *Agriosomyces*, *Ghazallomyces*, and *Khyollomyces*^50^). These results clearly demonstrate that the scope of AGF diversity is much broader than implied from prior studies. This conclusion is in apparent disagreement with the recent work of Paul et al.^17^, where the authors utilized a rarefaction-based approach on publicly available ITS1 AGF sequence data to suggest that AGF sampling efforts have reached saturation. However, we argue that using a rarefaction curve approach on publicly available datasets only elucidates coverage within samples already in the database, and not the broader AGF diversity in nature. Many prior studies have used relatively low throughput sequencing technologies, and repeatedly sampled few domesticated animals, and such pattern would result in encountering highly similar populations between different studies. We attribute the discovery and characterization of a wide range of novel AGF taxa within our dataset to sampling previously unsampled animal hosts, and the use of high-throughput sequencing that enabled access to rare members of the AGF community. Multiple novel AGF genera were isolated from animals previously unsampled for AGF diversity, e.g. *Aklioshbomyces* from white-tailed deer where it represented 98.5% of the community, *Ghazallomyces* from axis deer where it represented 27.8% of the community, and *Feramyces* from an aoudad sheep sample where it represented 55.3% of the community. It is notable that many of these novel taxa were only encountered in wild herbivores. Whether this novelty is a reflection of a lifestyle selecting for specific taxa, or a reflection of simply lack of prior sampling of wild animals due to logistic difficulties remains to be seen. This clearly demonstrates that novel AGF taxa remain to be discovered by sampling hitherto unsampled/poorly sampled animal hosts.

Further, a significant fraction of novel AGF candidate genera identified were present in extremely low relative abundance. Such pattern suggests the presence of numerous novel AGF taxa that appear to predominantly exist in relatively low abundance possibly as dormant members of the AGF community in the herbivorous gut. The discovery and characterization of the rare members of AGF community could significantly expand the scope of AGF diversity in nature. The dynamics, rationale for occurrence, mechanisms of maintenance, putative role in ecosystems, and evolutionary history of rare members of the community are currently unclear. It has been suggested that a fraction of the rare biosphere could act as a seed bank of functional redundancy that aids in ecosystem response to periodic (e.g. occurring as part of growth of the animal host, or due to seasonal changes in feed types) or occasional (i.e. due to unexpected disturbances) changes in the gut *in-situ* conditions. Regardless, such pattern highlights the value of deeper sampling (to capture rare biosphere), as well as more extensive time-series, rather than snapshot, sampling to capture patterns of promotion and demotion of members of the AGF community within the lifespan of an animal.

### The value of AGF isolation efforts

The strict anaerobic nature of AGF necessitates the implementation of special techniques for their isolation and maintenance^55, 56^. Further, while several storage methods based on cryopreservation have been proposed^57^, the decrease in temperature to the ultra-low values and the incidental O2 exposure during revival of the cryopreserved strains were shown before to be detrimental for some isolates. The lack of reliable long-term storage procedures often necessitates frequent subculturing of strains (every 3-4 days), which often leads to either the production of sporangia that do not differentiate to zoospores, or the outright failure to produce sporangia^58^.

Through a multi-year effort, we were successful in obtaining 216 isolates representing twelve AGF genera. We attribute our success to using multiple strategies (enrichment on multiple carbon sources, and paying special attention to picking colonies of different shapes and sizes, and to picking several colonies of the same shape, as representatives of different genera are known to produce colonies with very similar macroscopic features), but, more importantly, to using a wide range of samples (with varying host lifestyle, gut type, and animal phylogeny). The success of isolation of a certain genus was, in general, attributed to its abundance in the sample (Pearson correlation coefficient=0.79), especially for monocentric genera (e.g. *Pecoramyces*, *Feramyces*, *Neocallimastix*, and *Agriosomyces*), and was negatively correlated to the sample evenness (Pearson correlation coefficient= −0.87). It remains to be seen if this is true and reproducible for all samples and across laboratories. More efforts are certainly needed to develop targeted isolation strategies for specific taxa that we failed to obtain in pure cultures despite our best effort and despite their abundance in their respective sample (e.g. SK4 in one of the aoudad sheep samples, and RH5 in the domesticated sheep and the axis deer samples).

In conclusion, our results establish the utility of D1/D2 LSU and PacBio sequencing for AGF diversity surveys, and the culturability of a wide range of AGF taxa, and demonstrate that wild herbivores represent a yet-untapped reservoir of AGF diversity.

## Experimental Procedures

### Samples

Fecal Samples were obtained from six domesticated, six zoo-housed, and nine wild animals (Table 2). The host animals belonged to the families *Bovidae* (11), *Cervidae* (6), *Equidae* (3), and *Camelidae* (1). The dataset encompassed some replicates from few animal species sometimes with lifestyle variations within a single animal species: *Bos taurus* (n=2; domesticated cow, and domesticated longhorn cattle), *Ovis aries* (n=2; domesticated sheep and wild mouflon ram), *Capra aegagrus* (n=3; domesticated goat, wild Boer goat, and zoo-housed dwarf goat), and *Ammotragus lervia* (n=2; Aoudad Sheep) (Table 2). Samples from domesticated animals were obtained from Oklahoma State University and surrounding farms between September 2016 and May 2018. Samples from the Oklahoma City Zoo were obtained in April 2019. For samples from wild herbivores, we enlisted the help of hunters in four separate hunting expeditions in Sutton, Val Verde, and Coke counties, Texas (April 2017, July 2017, and April 2018), and Payne County, Oklahoma (October 2017). Appropriate hunting licenses were obtained and the animals were shot either on private land with the owner’s consent or on public land during the hunting season. All samples were stored on ice and promptly (within 20 minutes for domesticated samples, within 1 hour for zoo samples, and within 24 hours for samples obtained during hunting trips) transferred to the laboratory. Upon arrival, a portion of the sample was immediately used for setting enrichments for isolation efforts, while the rest was stored at −20°C for DNA extraction.

### Development of D1/D2 LSU locus as a phylogenetic marker

#### A. Amplification of the ITS1-5.8S rRNA-ITS2-D1/D2 LSU from Neocallimastigomycota isolates

Biomass was harvested from 10 ml of 2-4-day old cultures and crushed in liquid nitrogen. DNA was extracted from the ground fungal biomass using DNeasy PowerPlant Pro Kit (Qiagen, Germantown, Maryland) according to the manufacturer’s instructions (Youssef et al. 2013). A PCR reaction targeting the region encompassing ITS1, 5.8S rRNA, ITS2, and D1/D2 region of the LSU rRNA’ (Figure 6) was conducted using the primers ITS5-NL4^23^. The PCR protocol consisted of an initial denaturation for 5 min at 95 °C followed by 40 cycles of denaturation at 95 °C for 1 min, annealing at 55 °C for 1 min and elongation at 72 °C for 2 min, and a final extension of 72 °C for 20 min. PCR amplicons were purified using PureLink^®^ PCR cleanup kit (Life Technologies, Carlsbad, California), followed by cloning into a TOPO-TA cloning vector according to the manufacturer’s instructions (Life Technologies, Carlsbad, California). Clones (n=1-12 per isolate) were Sanger sequenced at the Oklahoma State University DNA sequencing core facility.

**Figure 6.**
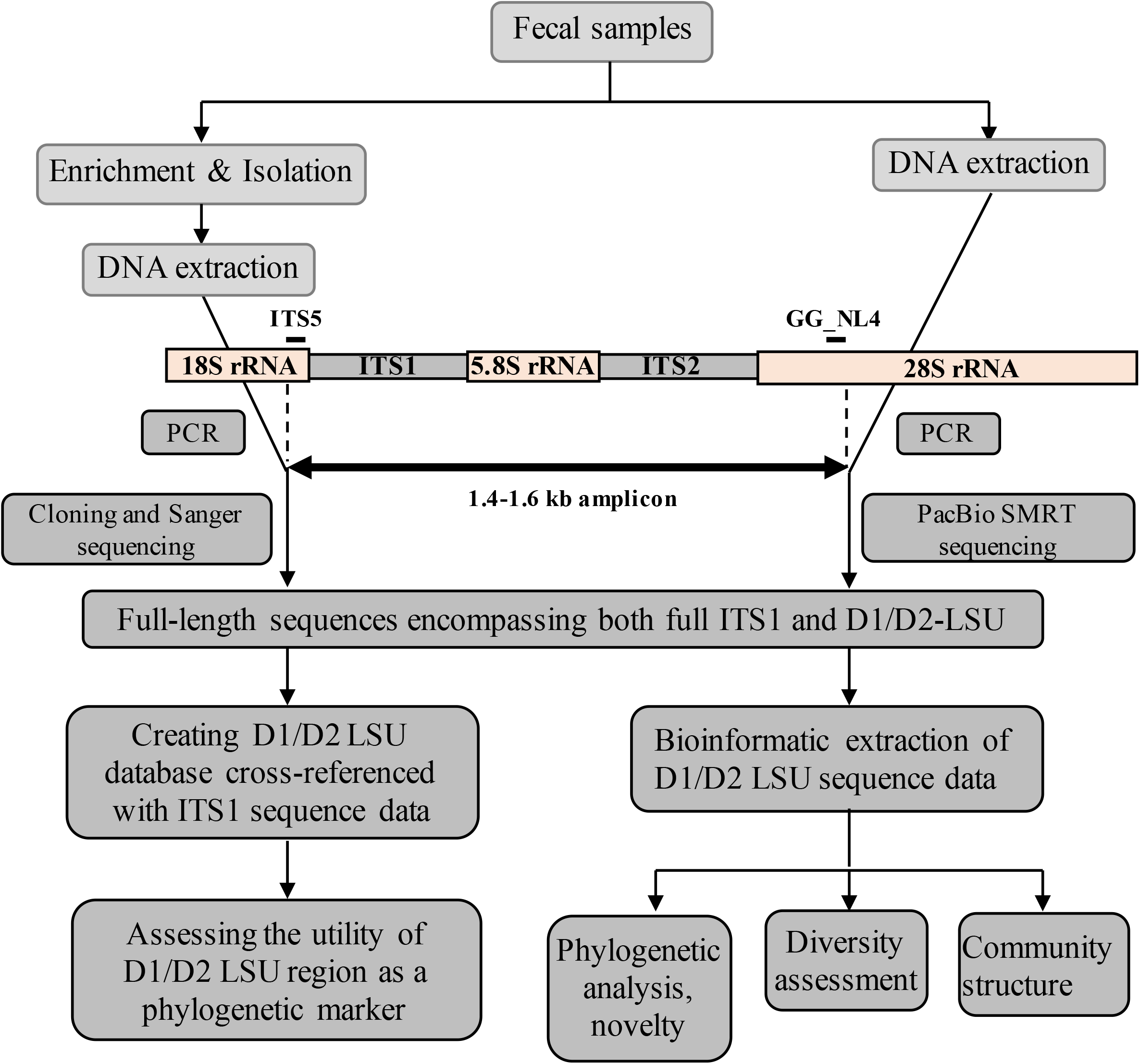
Workflow diagram describing the methods employed in this study.

#### B. Amplification of the ITS1-5.8S-ITS2-D1/D2 LSU from environmental samples

Fecal material from different animals (0.25-0.5 g) were crushed in liquid nitrogen and total DNA was extracted from the ground sample using DNeasy PowerPlant Pro Kit (Qiagen, Germantown, Maryland) according to the manufacturer’s instructions (Youssef et al. 2013). Extracted DNA was then used as a template for ITS1-5.8S-ITS2-D1/D2 LSU PCR amplification using ITS5 forward primer and the AGF-specific reverse primer GG-NL4^24^. Primers were barcoded to allow PacBio sequencing and multiplexing (Table S2). Amplicons were purified using PureLink^®^ PCR cleanup kit (Life Technologies, Carlsbad, California), quantified using Qubit^®^ (Life Technologies, Carlsbad, California), pooled, and sequenced at Washington State University core facility using one cell of the single molecular real time (SMRT) Pacific Biosciences (PacBio) RSII system.

#### C. Environmental PacBio-generated sequences quality control

We performed a two-tier quality control protocol on the generated sequences. First, the raw reads were processed according to PacBio published protocols to obtain single molecule consensus reads. Second, we used rigorous sequence quality control in Mothur^59^ to remove any sequences with low quality from subsequent analysis.

For Raw reads processing, the official PacBio pipeline (RS_Subreads.1) (http://files.pacb.com/software/smrtanalysis/2.2.0/doc/smrtportal/help/!SSL!/Webhelp/CS_Prot_RS_Subreads.htm) was utilized. Raw reads were filtered based on the minimum read length and minimum read quality. The passing reads were then processed with the PacBio RS_ReadsOfInsert protocol (http://files.pacb.com/software/smrtanalysis/2.2.0/doc/smrtportal/help/!SSL!/Webhelp/CS_Prot_RS_ReadsOfInsert.htm) for generating single-molecule consensus reads from the insert template. Consensus reads had a minimum of 5 full passes, 99.95% predicted accuracy, and 1000 bp insert length. The resulting consensus reads had a mean number of passes of 20, mean read length of insert of 1429 bp, and mean polymerase read quality of 0.99.

Sequence quality control procedures were subsequently conducted in Mothur^59^ to assess the quality of the generated consensus reads utilizing stringent protocols previously suggested for assessing bacterial, archaeal, and fungal diversity for similar sized amplicons^33, 52, 53, 54^. Reads were filtered in Mothur using trim.seqs to remove reads longer than 2000 bp, reads with average quality score below 25, reads with ambiguous bases, reads not containing the correct barcode sequence, reads with more than 2 bp difference in the primer sequence, and reads with homopolymer stretches longer than 12 bp. Reads with the primer sequence in the middle were identified by performing a standalone Blastn-short using the primer sequence as the query, and were subsequently removed using the remove.seqs command in Mothur.

A mock community (constituted of equal concentration of PCR products of 5 different strains (*Aklioshbomyces papillarum* strain WT2, *Feramyces austinii* isolate DS10, *Liebetanzomyces* sp. isolate Cel1A, *Piromyces* sp. isolate A1, and *Piromyces* sp. isolate Jen1) from our culture collection and for which we have obtained at least 5 Sanger clone sequences) was also sequenced. To establish whether the above approaches for overall read- and sequence-based quality control are adequate, we compared PacBio-generated mock sequences to the corresponding Sanger-generated clone sequences. The median percentage similarity between PacBio-generated sequences affiliated with a certain strain and the Sanger-generated clone sequences obtained for that strain (99.05±0.6 to 99.64±0.47) were not significantly different from the median percentage similarities between different clones of the same strain (98.91±0.6 to 99.72±0.47) (Student t-test p-value>0.1) attesting to the adequacy of the above quality control measures in removing low quality sequences.

#### D. A D1/D2 LSU reference database for cultured and yet-uncultured AGF taxa

A reference D1/D2-LSU sequence database for all Neocallimastigomycota cultured genera present in our culture collection was created via amplification, cloning, and sequencing of the ITS1-5.8S-ITS2-D1/D2 LSU allowing for a direct correlation and cross-referencing of both regions. To obtain D1/D2 LSU sequences representing yet-uncultured candidate genera previously defined by ITS1 sequence data^3, 5, 13, 17, 46^, the ITS1 region from the PacBio-generated ITS1-5.8S-ITS2-D1/D2 LSU environmental amplicons was extracted in Mothur using the pcr.seqs command with the sequence of the MNGM2 reverse primer and the flag rdiffs=2 to allow for 2 differences in primer sequence. The trimmed sequences corresponding to the ITS1 region were compared, using blastn, to a manually curated Neocallimastigomycota ITS1 database encompassing all known cultured genera, as well as yet-uncultured taxa previously identified in culture-independent studies^3, 5, 13, 17, 46^ (Figure 6). Sequences were classified as their first hit taxonomy if the percentage similarity to the first hit was >96% and the two sequences were aligned over >70% of the query sequence length. A taxonomy file was then created that contained the name of each sequence in the PacBio-generated environmental dataset and its corresponding taxonomy and was used for assigning taxonomy to the D1/D2 LSU sequence data.

#### E. Comparison of D1/D2-LSU versus ITS1 as phylogenetic markers

We used the dataset of full length PacBio-generated sequences described above, in addition to 116 Sanger-generated clone sequences from this study and previous studies^23, 28, 29, 49^, as well as genomic rRNA loci from two Neocallimastigomycota genomes (*Pecoramyces ruminantium* strain C1A, and *Neocallimastix californiae* strain G1)^36, 43^ in which the entire rrn operon sequence data is available to compare the ITS1 and D1/D2-LSU regions with respect to heterogeneity in length and intra-genus sequence divergence. For every sequence, the ITS1, and the D1/D2-LSU regions were bioinformatically extracted in Mothur using the pcr.seqs command (with the reverse primer MNGM2, and the forward primer NL1, for the ITS1, and the D1/D2-LSU regions, respectively) and allowing for two differences in the primer sequence. The trimmed sequences (both ITS1 and D1/D2-LSU) were then sorted into files based on their taxonomy such that for each genus/taxon two fasta files were created, an ITS1 and a D1/D2-LSU. These fasta files were then used to compare length heterogeneity, and intra-genus sequence divergence as follows. Sequences lengths in each fasta file were obtained using the summary.seqs command in Mothur. Intra-genus sequence divergence values were obtained by first creating a multiple sequence alignment using the MAFFT aligner^60^, followed by generating a sequence divergence distance matrix using the dist.seqs command in Mothur. Box plots for the distribution of length and sequence divergence were created in R.

### AGF Diversity assessment using D1/D2 LSU locus

#### A. Phylogenetic placement

The majority of the D1/D2-LSU sequences bioinformatically extracted from environmental sequences were assigned to an AGF genus as described above. D1/D2-LSU sequences that could not be confidently assigned to an AGF genus were sequentially inserted into a reference LSU tree to assess novelty. Further, the associated ITS1 sequences (obtained from the same amplicon) were similarly inserted into a reference ITS1 tree for confirmation. Sequences were assigned to a novel candidate genus when both loci (LSU and ITS1) cluster as novel, independent genus-level clades with high (>70%) bootstrap support in both trees.

#### B. Genus and species level delineation

Genus level assignments were conducted via a combination of similarity search and phylogenetic placement as described above. We chose not to depend on sequence divergence cutoffs for OTU delineation at the genus level since some genera exhibit high sequence similarity between their D1/D2-LSU sequences (e.g. *Liebetanzomyces, Capellomyces*, and *Anaeromyces* D1/D2-LSU sequence divergence ranges between 1.8-2.5%), while other genera are highly divergent (e.g. *Piromyces* intra-genus sequence divergence cutoff of the D1/D2-LSU region ranges between 0-5.7%), and as such “a one size fits all” approach should not be applied. On the other hand, a similar approach for OTUs delineation at the species equivalent level is problematic due to uncertainties in circumscribing species boundaries, and inadequate numbers of species representatives in many genera. Therefore, for OTU delineation at the species equivalent level, we reverted to using a sequence divergence cutoff. Historically, cutoffs of 3%^16^ to 5%^3^ were used for ITS1-based species equivalent delineation. However, D1/D2-LSU sequence data are more conserved when compared to LSU data in the Neocallimastigomycota^15, 26, 27, 28, 29, 30, 50^, as well as other fungi^19^. To obtain an appropriate species equivalent cutoff, we used the 116 Sanger-generated clone sequences from this study and previous studies^23, 28, 29, 49^, as well as genomic rRNA loci from two Neocallimastigomycota genomes where the entire ribosomal operon sequence is available (*Pecoramyces ruminantium* strain C1A, and *Neocallimastix californiae* strain G1)^36, 43^. The ITS1 and D1/D2-LSU regions were bioinformatically extracted and sorted to separate fasta files. Sequences in each file were then aligned using MAFFT^60^ and the alignment was used to create a distance matrix for every possible pairwise comparison using dist.seqs command in Mothur. The obtained pairwise distances for the ITS1, and the D1/D2-LSU regions were then correlated to obtain values of D1/D2-LSU sequence divergence cutoffs corresponding to the 3-5% range in ITS1. This was equivalent to 2.0-2.2%, and hence, for this study, a cutoff of 2% was used for OTU generation at the species equivalent level using the D1/D2-LSU region.

#### C. Diversity and community structure assessments

Genus and species equivalent OTUs generated as described above were used to calculate alpha diversity indices (Chao and Ace richness estimates, Shannon diversity index, Simpson evenness index) for the different samples studied using the summary.seqs command in Mothur. A shared OTUs file created in Mothur using the make.shared command was used to calculate Bray-Curtis beta diversity indices between different samples (using the summary.shared command in Mothur). The shared OTUs file was also used as a starting point for ranking the samples based on their diversity using both an information-related diversity ordering method (Renyi generalized entropy), and an expected number of species-related diversity ordering method (Hulbert family of diversity indices) (Table 2). For community structure visualization, Bray Curtis indices at the genus level were also used to perform non-metric multidimensional scaling using the metaMDS function in the Vegan package in R. Also, the percentage abundance of different genera across the samples studied were used in principal components analysis using the prcomp function in the labdsv package in R. Ordination plots were generated from the two analyses (NMDS and PCA) using the ordiplot function.

#### D. Statistical analysis

Correlations of the diversity estimates to animal host factors including the animal lifestyle (domesticated, zoo-housed, wild), and the animal host families (Bovidae, Cervidea, Equidae, Camelidae) were calculated using χ^2^-contingency tables followed by measuring the degree of association using Cramer’s V statistics as detailed before^3^. In addition, to identify factors impacting AGF diversity, Student t-tests were used to identify significant differences in the above alpha diversity estimates based on animal lifestyle (zoo-housed, domesticated, wild), and host phylogeny (families Bovidae, Equidae, Cervidae). To test the effect of the above host factors on the AGF community structure, Student t-tests were used to identify significant associations between specific AGF taxa and animal lifestyle (zoo-housed, domesticated, wild) or host family (families Bovidae, Equidae, Cervidae).

### Isolation efforts

Fecal samples (either freshly obtained, or stored at −20°C in sterile, air-tight plastic tubes) were used for isolation. Care was taken to avoid sample repeated freezing and thawing. Samples were first enriched by incubation for 24 h at 39°C in rumen-fluid-cellobiose (RFC) medium supplemented with antibiotics (50 μg/mL kanamycin, 50 μg/mL penicillin, 20 μg/mL streptomycin, and 50 μg/ mL chloramphenicol)^27, 28, 29, 50, 51^. Subsequently, enrichments were serially diluted by adding approximately 1 ml of enriched samples to 9 mL of RF medium supplemented with 1% cellulose or a mixture of 0.5% switchgrass and 0.5% cellobiose. Since fungal hyphae and zoospores are usually attached to the coarse particulates in the enrichment, serial dilutions were conducted using Pasteur pipettes rather than syringes and needles, as the narrow bore of the needle prevented the fecal clumps from being transferred. Serial dilutions up to a 10^−5^ dilution were incubated at 39°C for 24–48 h. Dilutions showing visible signs of growth (clumping or floating plant materials, a change in the color of cellulose, and production of gas bubbles) were then used for the preparation of roll tubes^55, 56^ on RFC agar medium. At the same time, and as a backup strategy in case the roll tubes failed to produce visible colonies, the dilution tubes themselves were subcultured and transferred to fresh medium with the same carbon source. Single colonies on roll tubes were then picked into liquid RFC medium, and at least three rounds of tube rolling and colony picking were conducted to ensure purity of the obtained colonies. To maximize the chances of obtaining isolates belonging to different genera, special attention was given, not only to picking colonies of different shapes and sizes, but also to picking several colonies of the same shape, as representatives of different genera could produce colonies with very similar macroscopic features. Isolates were maintained by biweekly subculturing into RFC medium. For long-term storage, cultures were stored on agar medium according to the procedure described by Calkins et al.^51^.

### Data accession

Sanger-generated clone sequences from pure cultures were deposited in GenBank under accession numbers MT085665 - MT085741. SMRT-generated sequences were deposited at DDBJ/EMBL/GenBank under the Bioproject accession number PRJNA609702, Biosample accession numbers SAMN14258225, and Targeted Locus Study project accession KDVX00000000. The version described in this paper is the first version, KDVX01000000.

## Acknowledgements

This work has been funded by the NSF-DEB grant 1557102 to N.H.Y. and M.S.E.

