## Supplemental document for "Assessing anaerobic gut fungal (Neocalliamstigomycota) diversity using PacBio D1/D2 LSU rRNA amplicon sequencing and multi-year isolation"

Table S1. Correlation coefficients (as Cramer's V statistic) between the various species richness and diversity indices calculated for each of the animals studied and the corresponding animal family or animal lifestyle. Student t-test p-values are shown for the testing the significant difference between the various animal families or lifestyles and the diversity indices in the table header. Significant p-values are shown in boldface.

| Test statistic | Factor/ Comparison | Observed number of OTUs | | Chao | | Ace | | Simpson evenness | | Shannon | | Diversity ranking | |
| --- | --- | --- | --- | --- | --- | --- | --- | --- | --- | --- | --- | --- | --- |
|  |  | Sp. Eq.* | Genus | Sp. Eq. * | Genus | Sp. Eq. * | Genus | Sp. Eq. * | Genus | Sp. Eq. * | Genus | Sp. Eq. * | Genus |
| Cramer V | Family | 0.24 | 0.32 | 0.2 | 0.39 | 0.49 | 0.31 | 0.26 | 0.33 | 0.34 | 0.21 | 0.34 | 0.1 |
| Student t-test p-value | Bovidae | 0.241 | **0.035** | 0.209 | **0.042** | 0.515 | 0.405 | 0.562 | 0.095 | 0.350 | 0.909 | 0.506 | 0.242 |
|  | Cervidae | 0.299 | 0.124 | 0.143 | 0.115 | 0.073 | 0.448 | 0.478 | 0.778 | 0.486 | 0.360 | 0.123 | 0.475 |
|  | Equidae | 0.931 | 0.450 | 0.800 | 0.604 | 0.108 | 0.873 | 0.539 | 0.382 | 0.320 | 0.791 | 0.067 | 0.103 |
| Cramer V | Lifestyle | 0.49 | 0.37 | 0.35 | 0.39 | 0.2 | 0.37 | 0.33 | 0.4 | 0.41 | 0.25 | 0.36 | 0.17 |
| Student t-test p-value | Zoo | 0.186 | 0.180 | 0.356 | 0.893 | 0.636 | 0.822 | 0.944 | 0.748 | **0.046** | 0.271 | 0.746 | 0.553 |
|  | Wild | 0.067 | 0.099 | 0.107 | 0.344 | 0.581 | 0.237 | 0.559 | 0.313 | 0.166 | 0.751 | 0.586 | 0.746 |
|  | Domestic | 0.529 | 0.662 | 0.425 | 0.368 | 0.275 | 0.122 | 0.477 | 0.147 | 0.677 | 0.459 | 0.786 | 0.339 |

* Sp.Eq.: species equivalent refers to OTUs_0.02_

Table S2. Primers and barcodes used in this study*

| Sample | Forward primer barcode | Reverse primer barcode |
| --- | --- | --- |
| Alpaca | ACACGCATGACACACT | CTGCGTGCTCTACGAC |
| American bison | TACTAGAGTAGCACTC | CGCGCTCAGCTGATCG |
| American elk | TACTAGAGTAGCACTC | ATGATGTGCTACATCT |
| Aoudad sheep (1) | CTATACATGACTCTGC | ATGATGTGCTACATCT |
| Aoudad sheep (2) | TCAGACGATGCGTCAT | ATGATGTGCTACATCT |
| Axis deer | TCAGACGATGCGTCAT | GCGCACGCACTACAGA |
| Blackbuck deer | CTATACATGACTCTGC | ACAGTCTATACTGCTG |
| Boer goat | CTATACATGACTCTGC | CTGCGTGCTCTACGAC |
| Domestic cow | TGTGTATCAGTACATG | CTGCGTGCTCTACGAC |
| Domestic goat | ACACGCATGACACACT | ATGATGTGCTACATCT |
| Domestic horse | ACACGCATGACACACT | ACAGTCTATACTGCTG |
| Domestic sheep | TGTGTATCAGTACATG | GCGCGATACGATGACT |
| Dwarf goat | TACTAGAGTAGCACTC | ACAGTCTATACTGCTG |
| Fallow deer | TCAGACGATGCGTCAT | GCGCGATACGATGACT |
| Longhorn cattle | TGTGTATCAGTACATG | GCGCACGCACTACAGA |
| Miniature donkey | CTATACATGACTCTGC | GCGCACGCACTACAGA |
| Mouflon ram | CTATACATGACTCTGC | GCGCGATACGATGACT |
| Oryx | TCAGACGATGCGTCAT | CGCGCTCAGCTGATCG |
| Pere David's deer | TACTAGAGTAGCACTC | GCGCGATACGATGACT |
| White-tail deer | TCAGACGATGCGTCAT | CTGCGTGCTCTACGAC |
| Zebra | CTATACATGACTCTGC | CGCGCTCAGCTGATCG |

Forward barcodes were added to the forward primer ITS5 (5'-GGAAGTAAAAGTCGTAACAAGG-3' ). Reverse barcodes were added to the reverse primer GG_NL4 (5'-TCAACATCCTAAGCGTAGGTA-3').

Figure S1. Phylogenetic tree constructed using the ITS1 sequences of representatives of all AGF genera/candidate genera. Sequences were aligned using the MAFFT aligner and maximum likelihood tree was constructed in FastTree. Bootstrap values are based on 100 replicates and are shown for branches with >50% bootstrap support.

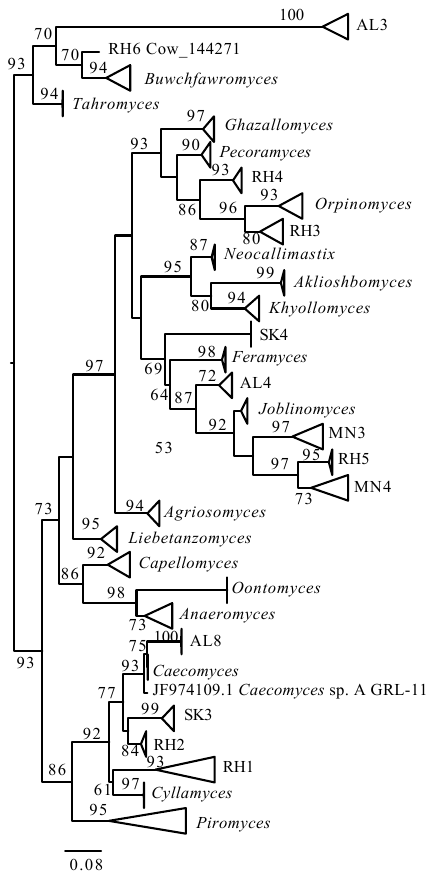

Figure S2. Principal component analysis plots based on percentage genus abundances in the different samples. Samples are shown as symbols and AGF genera affecting community structure are shown as '+'. Animals are displayed in black text while AGF genera are displayed in red text. (A) Symbols reflect lifestyle with domesticated animals shown as white squares, zoo-housed animals shown as grey squares, and wild animals shown as black squares. (B) Symbols reflect animal host phylogeny with Bovidae shown as squares, Cervidae shown as circles, Equidae shown as hexagons, and Camelidae shown as a star. Abbreviations: Am Bison, American bison; Ax deer, Axis deer; B goat, Boer goat; Bb deer, Blackbuck deer; Dw Goat, Dwarf goat; Fa deer, Fallow deer; Min Don, Miniature donkey; PD deer, Pere David's deer; WT deer, White-tail deer.

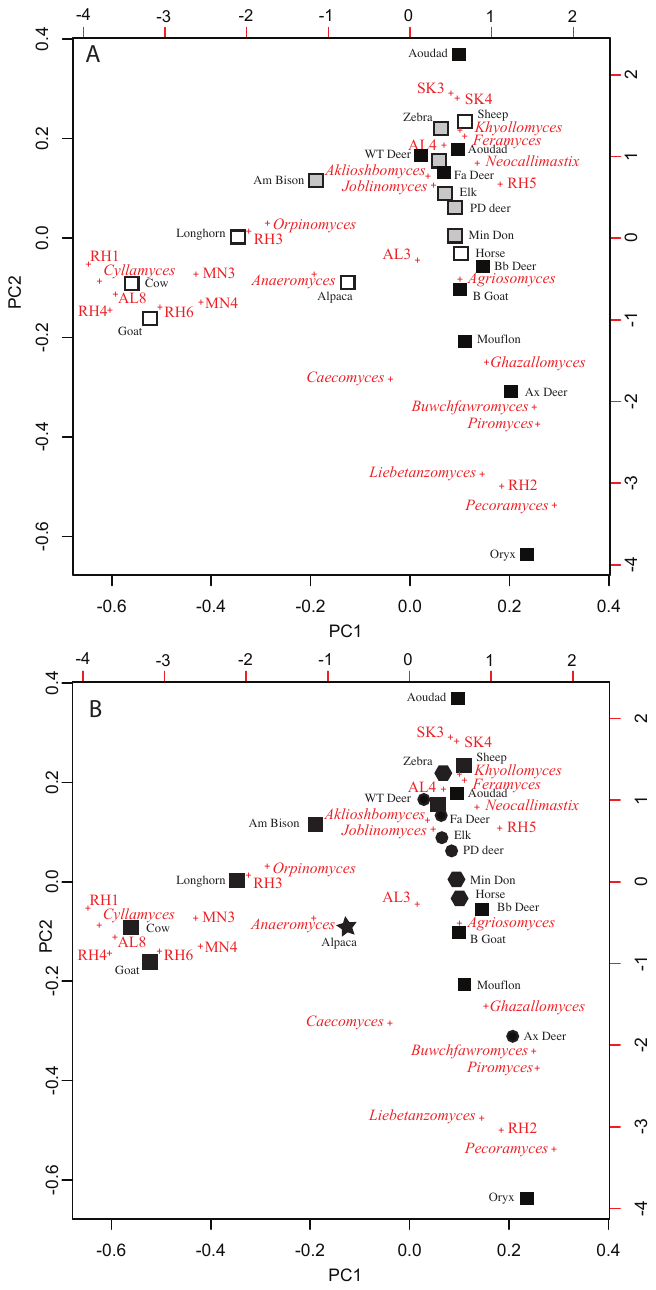

**Datasets 1-3.** Datasets generated in this study using SMRT amplicon sequencing. Each sequence name starts with the animal host and contains a unique identifier as well as the genus-level affiliation.

Dataset 1. Entire amplicon ITS1-5.8S rRNA-ITS2-D1/D2 LSU region (This dataset is also available in GenBank under Targeted Locus Study project accession KDVX00000000)

Dataset 2. ITS1 region bioinformatically extracted.

Dataset 3. D1/D2 LSU region bioinformatically extracted.
